# Gene drive dynamics in plants: the role of seedbanks

**DOI:** 10.1101/2025.04.24.649389

**Authors:** Isabel K. Kim, Leqi Tian, Ryan Chaffee, Benjamin C. Haller, Jackson Champer, Philipp W. Messer, Jaehee Kim

**Author notes:** These authors contributed equally (Isabel K. Kim, Leqi Tian). These authors jointly supervised this work (Jackson Champer, Philipp W. Messer, Jaehee Kim).

## Abstract

Gene drives offer revolutionary potential for the management of problematic plant populations, such as invasive weeds and herbicide-resistant species, by rapidly spreading desired genetic alterations. Two recent studies have provided the first experimental demonstrations of engineered CRISPR gene drive systems in plants (CAIN and ClvR). However, the successful application of such systems in the field will critically depend on an accurate understanding of plant-specific life-history traits, especially seed dormancy, a ubiquitous yet frequently overlooked eco-evolutionary force. In this study, we develop the first comprehensive modeling framework for gene drives in plant populations that incorporates a persistent soil seedbank. We show how the presence of a seedbank can significantly slow gene drive spread but also reduce the genetic load required to achieve population elimination. Furthermore, we show that seedbanks substantially increase the required introduction frequency of threshold-dependent gene drives, which could prevent establishment in some cases, yet also provide an intrinsic biosafety mechanism for confining a highly efficient drive to a target population. Our study highlights the need to incorporate seedbank dynamics into gene drive strategies to ensure realistic predictions and successful field applications.

## 1 Introduction

Weeds are among the most significant biological threats to global agriculture, causing annual yield losses in major crops that result in substantial economic damage and threaten food security [3–7]. Globally, weeds are responsible for around 10% of crop losses [3], and in the United States alone, they account for more than $26 billion a year in control expenses and lost crop yield [8]. The intensive and widespread use of chemical herbicides has historically provided the primary means of weed control; however, the rapid evolution of herbicide resistance poses an escalating challenge [9–13]. There is thus an urgent need to develop novel, effective, and evolutionarily robust weed management tools. One promising avenue for next-generation weed management is the use of gene drives—selfish genetic elements that can quickly propagate through populations, even if they confer a fitness cost to individual organisms [14–16]. Gene drives could be used either for population suppression or population modification of problematic weeds [17–22]. A suppression drive, for example, could spread deleterious traits (such as sterility or nonviability) to eradicate an invasive weed population, whereas a modification drive could reverse herbicide resistance by propagating susceptibility alleles. This raises the possibility that engineered gene drives could one day provide lasting, inexpensive solutions for controlling weeds that are otherwise difficult or costly to manage.

Recently, two toxin-antidote gene drive constructs were successfully developed and experimentally validated in the model plant *Arabidopsis thaliana* [23–25]: CRISPR-Assisted Inheritance using *NPG1* (CAIN; Liu et al. [1], Figure 1a) and Cleave-and-Rescue gamete killer (ClvR; Oberhofer et al. [2], Figure 1b). Unlike traditional gene drives that depend on homology-directed repair, an inefficient mechanism in plants due to their strong tendency towards end-joining DNA repair pathways [26], toxin-antidote gene drives avoid the need for homology-directed repair altogether. Instead, these drives spread by linking a toxin (Cas9 and guide RNAs targeting an essential gene) to an antidote (a tightly linked, cleavage-resistant copy of the target gene), resulting in elimination of genotypes carrying disrupted alleles but lacking the drive [27]. In experimental crosses, CAIN targeted *NPG1*, an essential gene required for pollen germination, achieving inheritance rates up to 97% through male gametes [1]. ClvR targeted *YKT61*, a gene required for viability in both pollen and ovules, achieving near-complete inheritance through males, and substantial, though lower, inheritance rates through females [2]. Preliminary modeling in both studies indicated that such drives could spread rapidly and reliably eliminate populations under panmictic, outcrossing conditions [1, 2].

**Figure 1.**
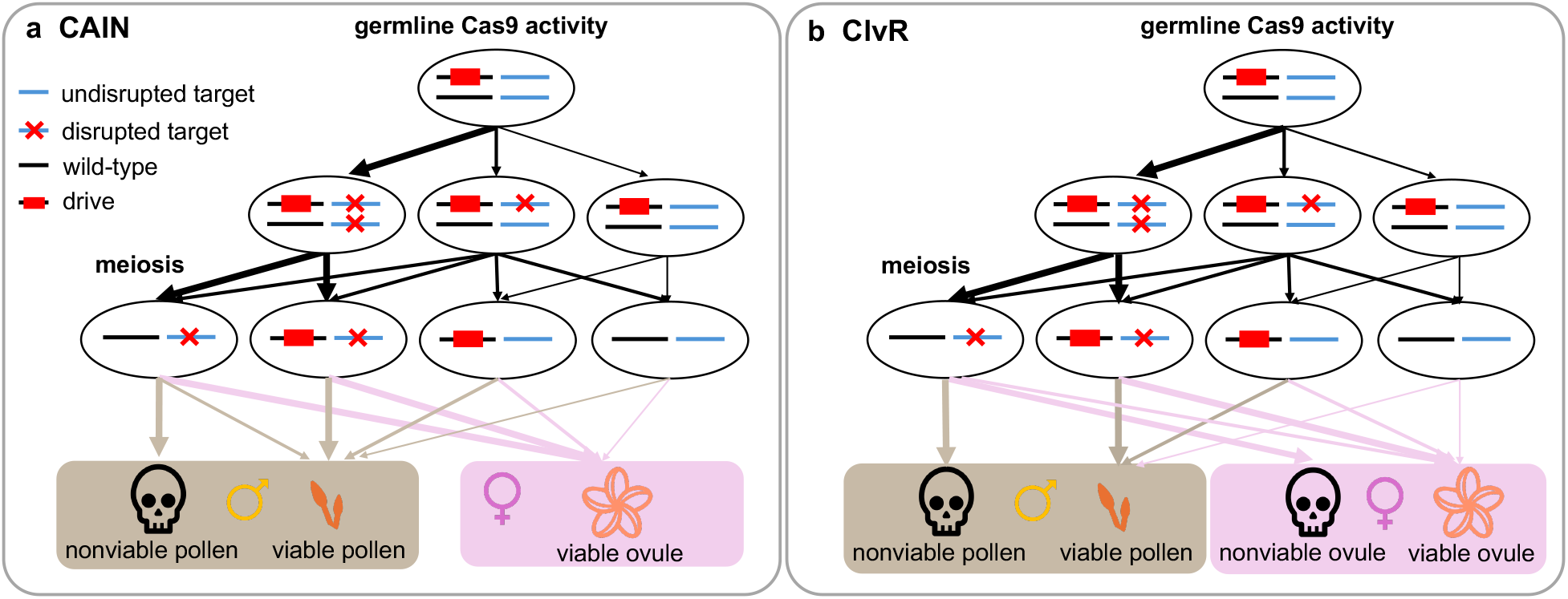
Schematic of gene drive systems. **a**, CAIN drive [1]. In the germline of a drive carrier, the drive cleaves each wild-type target allele at a rate equal to the sex-specific germline cleavage rate. The target gene is essential only for pollen germination; thus, ovules of any genotype are viable. Pollen inheriting the drive construct are always viable, since the drive contains a cleavage-resistant recoded functional target allele. A pollen grain with a wild-type allele and a disrupted target gene (leftmost gamete, bottom row) is nonviable with probability equal to the penetrance rate of the target gene (estimated at 96%). For suppression, we define the “CAIN male suppression” drive as the CAIN drive inserted into an essential haplosufficient male fertility gene, such that males with two drive alleles are sterile. **b**, ClvR drive [2]. ClvR functions similarly to CAIN but targets a fully penetrant gene that is essential for viability in both pollen and ovules. However, ovules inheriting disrupted alleles and no drive allele from drive-carrying females (leftmost gamete, bottom row) remain viable with probability equal to a maternal carryover rate (estimated at 20.7%); otherwise, they are nonviable. For suppression, we define two drives. “ClvR male suppression” targets an essential haplosufficient male fertility gene, such that homozygous males are sterile, and “ClvR female suppression” targets an essential haplosufficient female fertility gene, such that homozygous females are sterile.

However, these initial plant gene drive studies left out a key aspect of plant life history: seed dormancy. Dormancy allows seeds to remain in the soil without germinating for extended periods. This trait leads to a persistent seedbank, which can provide a genetically diverse reservoir of alleles in the soil that can buffer the population against rapid evolutionary or environmental changes [28–31]. In the presence of a seedbank, gene drive alleles might spread more slowly, extending the time to population modification or suppression [18, 32, 33]. Conventional models without seedbanks assume that every individual reproduces in each generation, so they cannot capture these age-structured, time-delayed dynamics. Yet, neither the CAIN nor ClvR modeling efforts [1, 2] included dormancy or seedbank processes, leaving a major gap in our understanding of gene drive behavior in real plant populations, especially given how common seedbanks are among weeds. There is thus a need to incorporate seedbank dynamics into gene drive models, not only to improve theoretical accuracy, but also to guide practical decisions about gene drive design, risk assessment, and field deployment. In particular, an understanding of seedbank effects should inform the choice of drive architecture, help determine how many modified individuals should be released, and set realistic expectations for the timeline and ultimate efficacy of a gene drive under field conditions.

In this study, we address this critical knowledge gap by developing a detailed, individual-based model for the eco-evolutionary dynamics of gene drives in plant populations that includes realistic life cycles and seedbank features. Our model focuses on dioecious, annual, diploid weeds species in which outcrossing is obligate and persistent seedbanks are commonly formed. We simulate both modification and suppression versions of CAIN and ClvR across a range of demographic and ecological conditions, varying seed germination rates and fecundity parameters. We specifically evaluate how seedbank characteristics affect (i) the fixation probability and time to fixation of the drive, (ii) the suppression potential of different drive constructs, and (iii) the introduction threshold frequency of the drive under different fitness costs.

## 2 Results

We developed a comprehensive simulation model of CAIN [1] and ClvR [2] (Figure 1), the only experimentally demonstrated gene drive constructs in plants, using the individual-based forward-time simulation framework SLiM v4.0.1 [34]. A detailed description of our model is provided in the Methods section, and all major symbols used in this work, along with their definitions and default values (where applicable), are summarized in Tables S1 and S2.

Briefly, each drive was modeled as either a modification drive (carrying a desirable allele) or a suppression drive (inducing sterility in drive homozygotes of one sex). Populations were initialized at carrying capacity (*K*) and allowed to equilibrate before introducing the drive in a single release. We explicitly modeled the life cycle of dioecious annual plants with seedbank dynamics and obligate outcrossing, assuming wind pollination and considering only the “effective” gametes that contribute to reproduction (Figure 2). Newly produced seeds entered the seedbank and experienced age-dependent survival and germination, yielding total germination probability *γ* and average seedbank duration *τ*. These metrics depend on the baseline survival rate (*d*, seed survival through the first year), baseline germination rate (*b*, first-year germination conditional on survival), and parameters governing age-dependent declines in seed survival (*q*) and germination (*m*). Seeds exceeding the maximum age *L* were removed from the seedbank.

**Figure 2.**
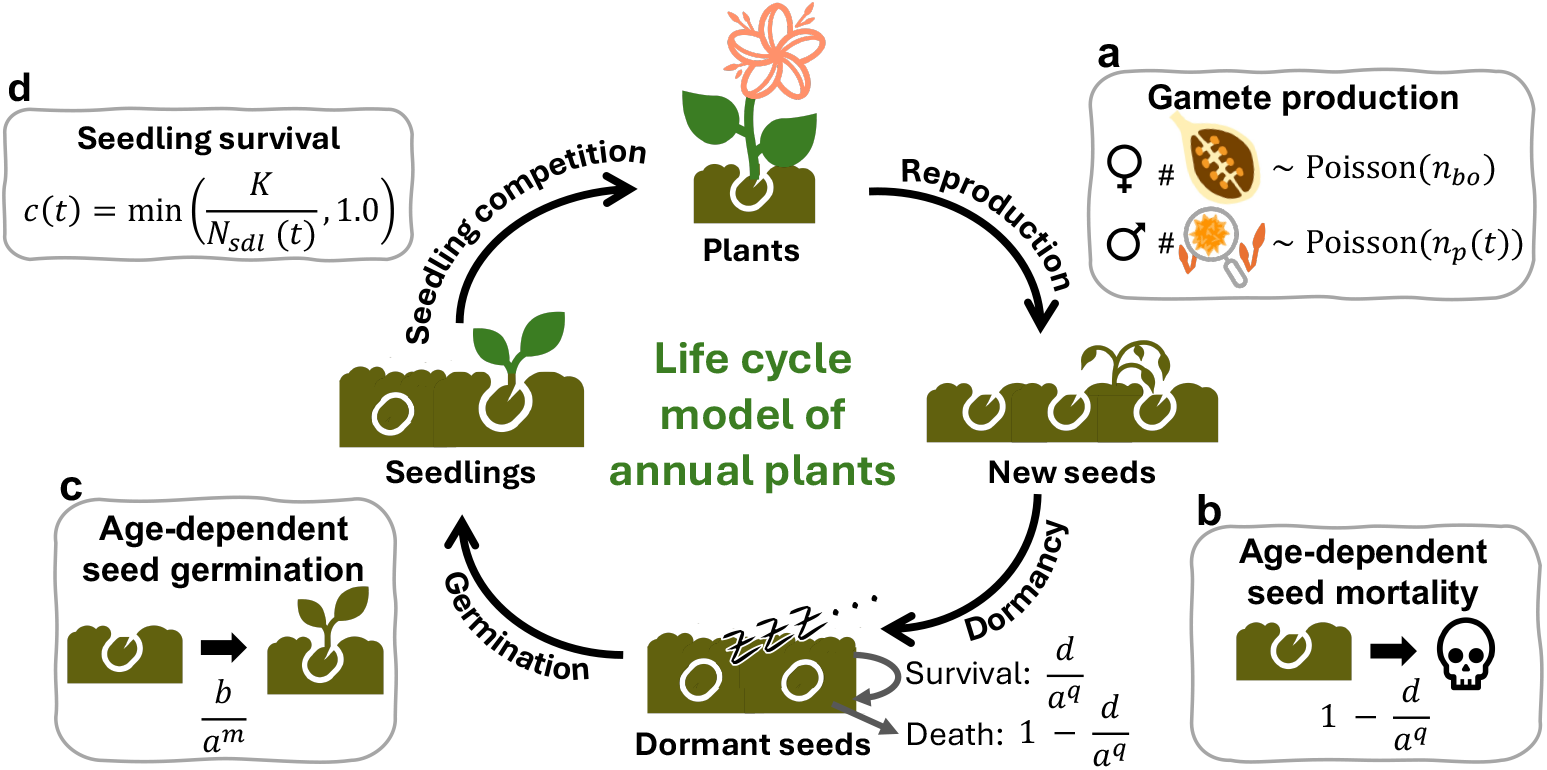
Life cycle model of annual plants. Illustration of the life cycle modeled for an dioecious annual plant population. **a**, Plants to seeds: an “effective ovule” is defined as an ovule capable of fertilization and subsequent seed development and an “effective pollen grain” as a pollen grain that can successfully reach a fertile female plant via wind and germinate. Each fertile female plant produces a Poisson-distributed number of effective ovules with mean *n*_*bo*_. Each fertile male plant produces a Poisson-distributed number of effective pollen grains with mean *n*_*p*_(*t*), which is proportional to the current number of fertile females. Effective pollen grains are randomly distributed among fertile female plants, and fertilized ovules develop into seeds. After reproduction, all plants are removed from the population. **b–c**, Into and out of the seedbank: age-dependent seed mortality and germination. A seed survives its first year at rate *d*, after which survival declines with seed age, such that a seed of age *a* survives at rate *d/a*^*q*^, where *q* modulates age-dependent mortality. Similarly, germination occurs initially at rate *b* and declines with seed age, such that a seed of age *a* germinates at rate *b/a*^*m*^, where *m* modulates age-dependent germination. **d**, Seed to plants: seedling competition. Germinated seeds become seedlings, which experience density-dependent competition before becoming adult plants. Each seedling at year *t* survives to adulthood with probability 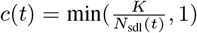, where *N*_sdl_(*t*) is the current number of seedlings, and *K* is the carrying capacity.

We analyzed drive outcomes across seedbank durations, total germination probabilities, and fecundities (characterized by the mean effective ovule count *n*_*bo*_ and mean effective pollen count *n*_*bp*_). Suppression drive efficacy was quantified by genetic load (*λ*), defined as the proportional reduction in effective gamete production relative to wild-type populations, with the required genetic load (*λ*^*\**^) denoting the threshold necessary for population elimination. Formal definitions of seedbank metrics and genetic load are provided in Eqs. 1–4.

### 2.1 Effect of a seedbank on modification drives

We found that both CAIN and ClvR performed effectively as modification drives over a wide range of seedbank parameters. When varying the baseline germination rate (*b*) from 0.05 to 1 and the age-dependent germination parameter (*m*) from 0 (age-independent germination) to 2 (germination rates strongly decreasing with age), both drives consistently reached fixation. In our model, setting *b* = 1 (with the default baseline survival rate *d* = 1) causes all seeds to survive and germinate immediately, eliminating dormancy beyond one year and making *m* irrelevant.

Though fixation was always achieved, the average time required for the drive to reach fixation from a 10% introduction frequency varied substantially across the parameter range (Figures 3 and S1) and was found to be positively correlated with the average seedbank duration *τ* (Figure S2a). Fixation occurred most rapidly when both *b* and *m* were high, conditions under which *τ* was low because germinating seeds were predominantly younger. Conversely, fixation was slowest under low baseline germination rates and minimal age-dependence, conditions under which *τ* was high, as a large proportion of germinating seeds were older. The relationship between *τ* and drive fixation time is intuitive, as *τ* determines the average generation time of the population, and gene drive spread is inherently a generational process. This result aligns with previous studies demonstrating that a seedbank slows the rate of selection [35, 36] and, similarly, the spread of gene drives [18, 32].

**Figure 3.**
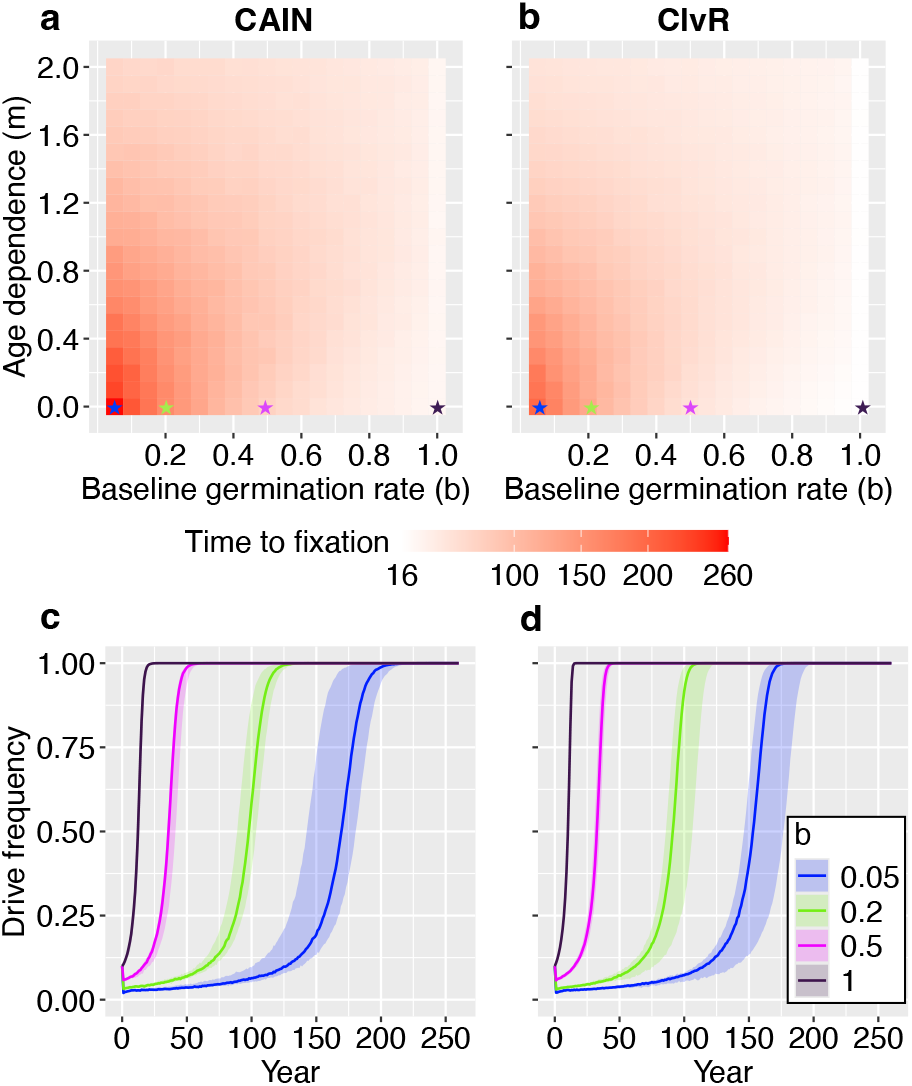
Spread of CAIN and ClvR modification drives under varying seedbank parameters. **a**, Mean fixation time for the CAIN drive across baseline germination rates (*b*) and age-dependent germination parameters (*m*); all other parameters were fixed at their default values (Table S2). Fixation is defined as 100% drive frequency in both plants and seeds. Colors represent mean fixation time across 10 replicates. CAIN achieved fixation under all tested parameter combinations. **b**, Same as **a** but for the ClvR drive, which also achieved fixation under all tested parameter combinations. **c**, CAIN drive frequency trajectories in plants for four baseline germination rates (*b*) with age-independent germination rates (*m* = 0). Parameter combinations correspond to stars of matching color in **a**. Solid lines represent median trajectories; shaded regions indicate the observed range (minimum–maximum) across 10 replicates. **d**, Same as **c** but for the ClvR drive, with parameter combinations corresponding to stars of matching color in **b**.

Across all parameter combinations we investigated, ClvR spread more rapidly than CAIN because ClvR biases inheritance in both sexes, while CAIN biases inheritance only through males. The disparity in fixation time between the two drives was most pronounced at higher values of *τ*. Figure S3 displays these time-to-fixation differences under default and low fecundity conditions. Changes in population fecundity did not affect the general patterns observed: both CAIN and ClvR still reached fixation across all germination rates, with time to fixation increasing with *τ* and ClvR consistently reaching fixation faster than CAIN (Figure S4).

### 2.2 Effect of a seedbank on suppression drives

To evaluate suppression drive performance across seedbank conditions, we varied the baseline germination rate (*b*) and the age-dependence germination parameter (*m*), while keeping drive-specific parameters fixed at their experimental values. We tracked the proportion of replicates in which the drive achieved population elimination and the mean elimination times following a 10% release in heterozygotes. We found that population elimination could not occur unless the drive reduced the effective ovule or pollen pool by a critical threshold, defined as the required genetic load (*λ*^*\**^). This threshold depends on the population total germination probability (*γ*) and mean effective ovule count (*n*_*bo*_) if ovules production is limited, or mean effective pollen count (*n*_*bp*_) if pollen production is limited (Figure 4 and Eq. 4). When individuals produce fewer effective gametes on average, the low-density growth rate of the population decreases, reducing the required drive strength. When the population has a low total germination probability, a large portion of seeds do not mature to adults, such that the population can collapse with lower genetic load from the drive. We defined the genetic load at time *t, λ*(*t*), as the proportional reduction in effective gamete pool due to the drive (Eq. 3) and examined which suppression drive types and conditions could achieve *λ*(*t*) *> λ*^*\**^ for population elimination.

**Figure 4.**
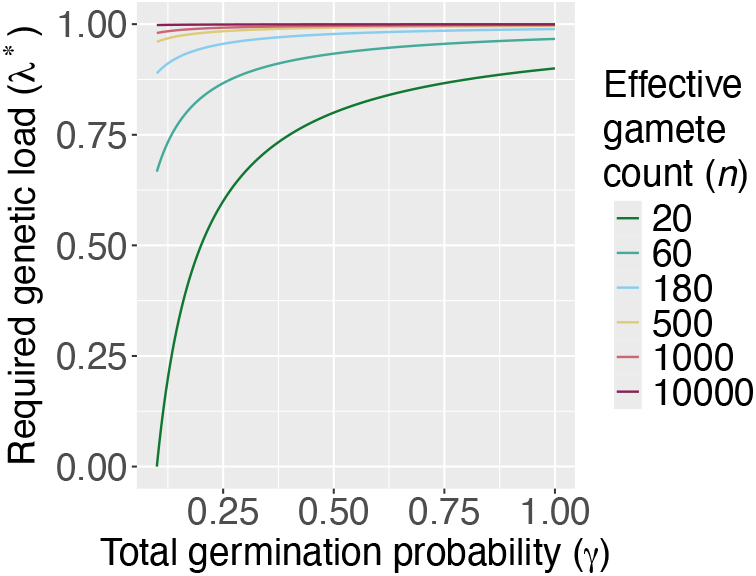
Required genetic load for population elimination via suppression drive. Genetic load is defined as the proportional reduction in the number of effective gametes due to the suppression drive (Eq. 3). The required genetic load (*λ*^*\**^) is a function of the total germination probability (*γ*) and the effective gamete count (*n*) of the sex being limited by the drive (Eq. 4). For male-targeting drives, *n* represents the mean number of effective pollen grains per male (*n*_*bp*_), where an effective pollen grain is one capable of reaching a receptive female and germinating. For female-targeting drives, *n* corresponds to the mean number of effective ovules per female (*n*_*bo*_), where an effective ovule is one that can be fertilized and develop into a seed.

The strongest suppression drive considered was the ClvR male suppression drive, in which drive homozygous males are sterile [2]. This drive benefits from high cleavage rates and a fully penetrant target gene that acts in both male and female gametes. It successfully achieved population elimination across all germination parameters tested (Figures S5a and S6a–c), with the average time to elimination (Figure 5a) being proportional to the average seedbank duration (*τ* ; Figure S2a). Figures 5b–d show the genetic load imposed by the drive over time. When there is no seedbank (i.e., *b* = 1), close to 99% of effective pollen must be removed from the gamete pool for the population to decline (Figure 5d). The ClvR male suppression drive can achieve this genetic load, causing a population crash once the required genetic load is reached (after about 33 years). At lower *b* (holding *m* constant), the total germination probability decreases (Figure S2b). As a result, fewer seeds survive dormancy, thus lowering the genetic load required for population elimination through pollen reduction (Figures 5b–c). While the drive can always reach this lower *λ*^*\**^, the increased generation time (*τ*) prolongs the time to suppression. When population fecundity is reduced, *λ*^*\**^ is lower across all *b* and *m* (Figure 4). The drive can therefore eliminate the population with even less pollen reduction (Figures S7a and S8a–c) and achieve slightly faster times to elimination (Figure S9a–d).

**Figure 5.**
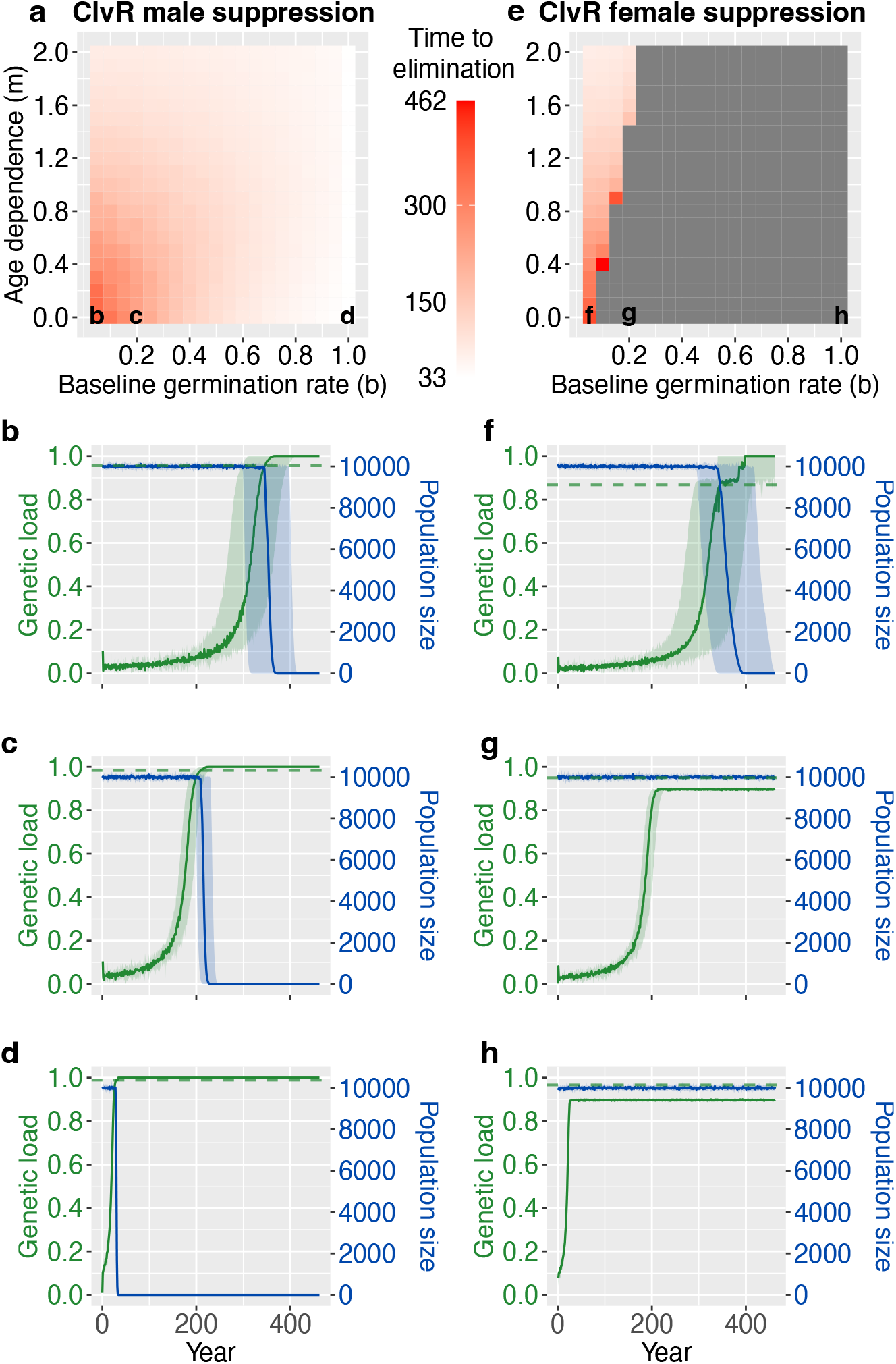
Dynamics of ClvR suppression drives under varying seedbank parameters. **a**, Heatmap of the mean time to population elimination for the ClvR male suppression drive across baseline germination rates (*b*) and age-dependent germination parameters (*m*); all other parameters were fixed at their default values (Table S2). Population elimination is defined as the complete loss of both seeds and plants due to the drive. The drive succeeded for all tested parameters (Figure S5a). Colors represent mean elimination time across 10 replicates; lettered cells correspond to subplots below. **b–d**, Trajectories of genetic load (green; left axis) and corresponding population size (blue; right axis) for ClvR male suppression drive, assuming age-independent germination rates (*m* = 0), with baseline germination rates: **b**, *b* = 0.05; **c**, *b* = 0.2; and **d**, *b* = 1. Genetic load is defined for this drive as the proportional reduction in total effective pollen grains in the population. Solid lines represent median trajectories; shaded regions indicate the observed range (minimum–maximum) across 10 replicates. The horizontal green dashed lines denote the required genetic load for population elimination (*λ*^*\**^; Eq. 4). **e**, Heatmap as in **a** but for the ClvR female suppression drive. The drive succeeded only for a subset of tested parameter sets (Figure S5b). Colors indicate mean elimination times among successful replicates; grey cells denote parameter sets where elimination consistently failed. Letter labels in the bottom row indicate parameter sets analyzed in subplots below. **f–h**, Same as **b**–**d** but for the ClvR female suppression drive, with genetic load defined as the proportional reduction in total effective ovules in the population.

When the ClvR drive targets a haplosufficient essential *female* fertility gene, its performance declines. Although this drive retains high cleavage rates and a fully penetrant target gene (as in the male suppression variant), drive inheritance is weaker through females because of maternal carryover: ovules with a disrupted target allele and no drive allele can remain viable if they inherit rescue protein from a drive-carrying mother [2]. Experimental results suggest this occurs at around a 20.7% rate (Section S1), allowing a greater fraction of female gametes to remain viable without a drive allele and reducing the drive frequency in the effective ovule pool relative to the effective pollen pool. Thus, the population can tolerate greater loss of drive alleles through male gametes than through female gametes, making sterile drive homozygous males a more effective suppression strategy than sterile drive homozygous females. This asymmetry is evident when comparing the parameter regimes for successful suppression in Figure S5.

Unlike the ClvR male suppression drive, the ClvR female suppression drive can eliminate the population only under very low total germination probabilities, specifically at low *b* and high *m*. In this parameter range, population elimination occurs more rapidly as *m* increases (Figure 5e), since the average seedbank duration decreases (Figure S2a). Figure 5h shows *λ*^*\**^ for a population without a seedbank, where for this drive, genetic load is defined as the relative reduction in the total number of effective ovules induced by the drive. In the absence of a seedbank, population decline requires the elimination of approximately 97% of the effective ovule pool. However, the ClvR female suppression drive cannot achieve this genetic load; instead, it spreads to a high equilibrium frequency, eliminating roughly 90% of effective ovules, but still permitting enough seed production to maintain the population (Figure S6d–f). When *b* is lowered to 0.2, the ClvR female suppression drive still failed to reach the genetic load required for population elimination (Figure 5g). Only when *b* is further reduced to 0.05 can the drive reach the necessary genetic load (Figure 5f). Under these conditions, approximately 75% (1 *− γ*) of seeds fail to germinate from the seedbank, reducing the required effective ovule pool elimination to about 87%, a level lower than the drive’s maximum attainable genetic load. The drive reaches the required genetic load around year 345 (i.e., generation 115.7), after which point, the population declines. When the population’s fecundity is reduced, the required genetic load (*λ*^*\**^) decreases across all *b* and *m* (Figure S9f–h). As a result, the drive can eliminate the population across a wider range of seedbank parameters (Figures S7b, S8d–f, and S9e).

The CAIN male suppression drive (where drive homozygous males are sterile; [1]) behaves similarly to the ClvR female suppression drive, lacking sufficient suppressive power to eliminate populations with high *γ* (Figure S10a). This drive spreads exclusively through males and targets a gene with incomplete penetrance. With *NPG1* penetrance estimated at 96%, approximately 4% of pollen grains carrying disrupted target alleles remain viable without a drive allele. Thus, unlike ClvR, CAIN does not bias inheritance through females and exhibits slightly weaker inheritance bias through males. This, combined with the loss of drive alleles via sterile drive-homozygous males, restricts CAIN’s ability to eliminate the population to scenarios where the seedbank removes a large fraction of seeds (Figures S10a and S2b) or fecundity is low (Figure S10b). In the parameter regime allowing suppression, the time to population elimination is shortest at higher *m*, where *τ* is lower (Figures S11a and e). Figure S11d shows the equilibrium genetic load of the drive when there is no seedbank. At our default population fecundity, the drive must remove about 99% of effective pollen to achieve population elimination, but its inefficiencies cap the genetic load at 96%. As a result, the drive attains a high equilibrium frequency yet fails to remove enough effective pollen to cause population decline (Figure S12a–c). When the baseline germination rate is reduced to *b* = 0.2 (with *m* = 0), the required genetic load is 98%, still exceeding the maximum achievable genetic load by the drive (Figure S11c). Reducing *b* to 0.05 lowers the required load to 96%, enabling population elimination (Figure S11b). Likewise, under reduced fecundity, the drive can attain the required genetic load across a broader range of *b* and *m* (Figures S10b, S11e–h, and S12d–f).

### 2.3 Effect of a seedbank on threshold-dependent gene drives

So far, we have assumed that neither CAIN nor ClvR incur a fitness cost, making them effectively zero-threshold drives. Though spread is slow until an intermediate drive frequency is reached, either drive will spread as long as it escapes stochastic loss when still at low frequencies. However, when fitness costs are present, these drives become threshold-dependent, requiring an introduction above their invasion threshold frequency to ensure spread [2, 27]. For example, a fitness cost reducing gamete viability (*s*_*g*_ *>* 0) lowers average seed production in drive carriers relative to wild-type individuals, thereby impeding drive invasion at low frequencies. Alternatively, a fitness cost reducing seed survival (*s*_*s*_ *>* 0) results in wild-type seeds having higher probabilities of survival and germination compared to drive-carrying seeds. To overcome this disadvantage in the seedbank, a higher initial introduction frequency of drive individuals is necessary. In this section, we explore the consequences of fitness costs on the invasion threshold of the drive.

We first examined the effects of a fitness cost reducing gamete viability *s*_*g*_ *>* 0 with *s*_*s*_ = 0. Here, *s*_*g*_ defines the probability that a drive-carrying gamete is nonviable, acting as a codominant fitness cost on fecundity. Under this model, drive homozygotes are expected to produce half as many viable gametes as heterozygotes. We define 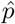 as the baseline invasion threshold of the drive in the absence of a seedbank (i.e., with *b* = 1) and 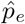 as the effective invasion threshold of the drive in the presence of a seedbank. For simplicity, we consider only a single release of the drive, rather than a sustained multi-year release.

Because some seeds die or fail to germinate, we expect 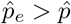, with the extent of this difference being dependent on seedbank parameters. To estimate 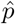 for a given drive and fitness cost, we varied the drive’s introduction frequency (*p*_0_) and identified the lowest frequency above which the drive spread in more than 50% of replicates. To fully explore introduction frequencies (including *p*_0_ *>* 0.5), we introduced the drive by converting *p*_0_ of plants to drive homozygotes, in contrast to previous sections, where the drive was introduced exclusively in heterozygotes. We decreased the baseline germination rate *b* and estimated the effective invasion threshold 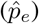 using the same procedure as above. The results (Figures S13 and S14) suggest a roughly proportional relationship between the effective invasion threshold and the average seedbank duration: 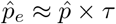.

When the average seedbank duration (*τ*) is low, younger seeds are more likely to germinate than older ones; hence, the drive can be introduced at a frequency closer to 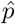, as many drive seeds germinate soon after release. However, when *τ* is high, this process experiences a greater temporal lag, necessitating a higher introduction frequency. In the initial years following drive release, most germinated seeds originate from a period when only wild-types were present, causing an initial decline in the drive frequency among plants. However, over time, as seeds produced after the release begin to germinate, the fraction of drive-carrying germinated seeds increases. Once the influx of drive-carrying seeds causes the drive frequency to exceed 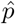 in the plant population, the drive is set on a trajectory toward fixation, expected to rise in frequency each subsequent year, with *τ* capturing this temporal delay. Furthermore, a longer average seedbank duration requires a higher initial drive frequency to offset the initial influx of older wild-type seeds and ensure eventual drive establishment.

One major implication of this relationship is that for long-lived seedbanks (high *τ*) or drives with high baseline invasion thresholds, 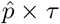 can exceed 1. In such cases, even releasing the drive at 100% frequency in plants cannot prevent its eventual loss (e.g., Figures S13k–l and S14k–l). Shortly after release, germination is predominantly from older wild-type seeds, pushing the drive frequency in plants below 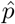. While a transient drop in drive frequency does not guarantee drive loss, germinated drive seeds must ultimately push the drive frequency in plants above 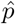 for the drive to spread. However, when 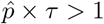, even releasing the drive at the maximum possible frequency in plants cannot offset the dilution effect imposed by the seedbank. Thus, with a single release, the drive can never spread above 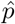 in the plant population, ultimately leading to the drive’s loss.

Figure 6 compares 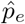 predicted by 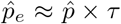 with the observed 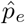 for both modification drives. At lower fitness costs, this equation accurately predicts the threshold; however, at the highest fitness cost explored (*s*_*g*_ = 0.25), this equation slightly underestimates the observed value. These trends also hold for suppression drives (Figure S15). For the suppression drives causing male sterility, we introduced the drive by converting a fraction *p*_0_ of the population to homozygous females; for the suppression drive causing female sterility, we converted *p*_0_ of the population to homozygous males. Given an equal sex ratio, the maximum achievable *p*_0_ in either case was 0.5. We found that if 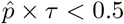, the prediction holds, and if 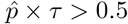, the suppression drive cannot spread (Figures S15–S18).

**Figure 6.**
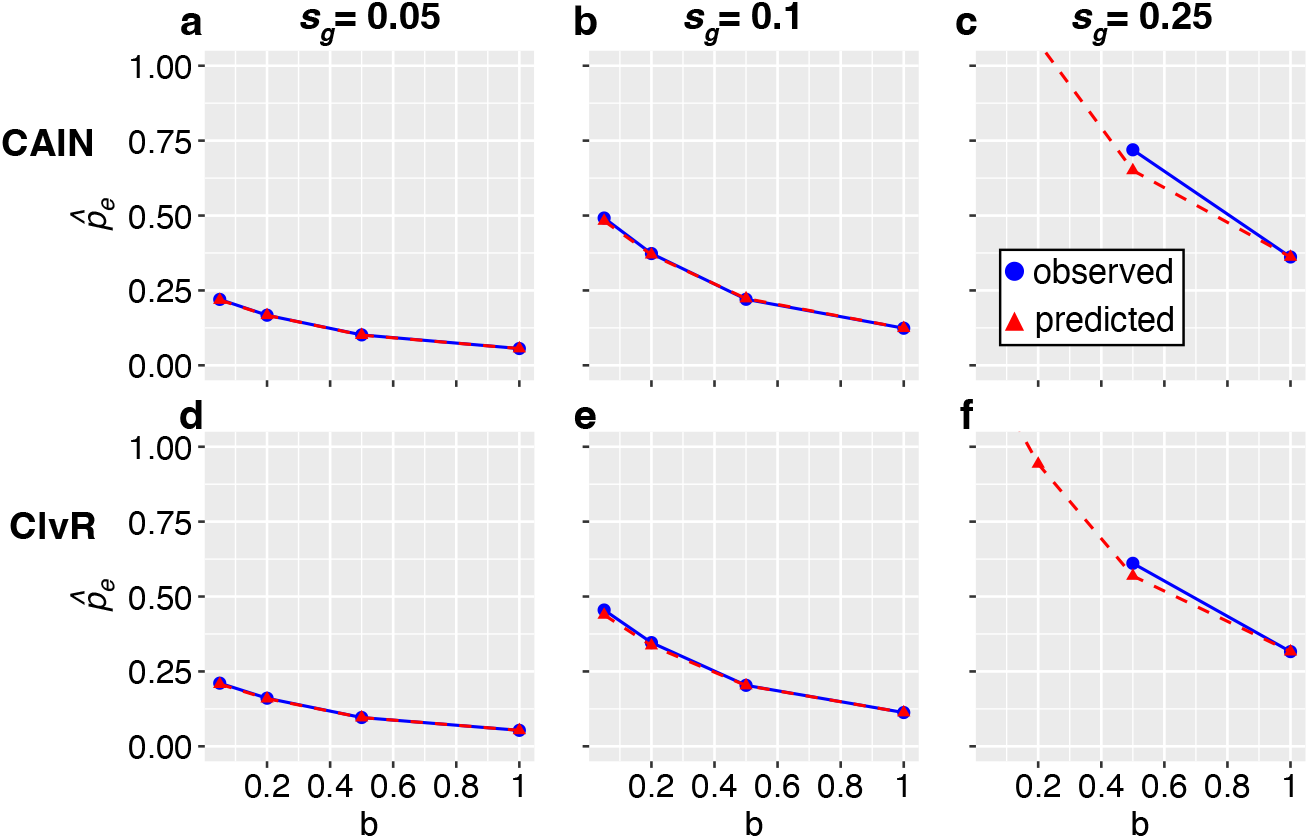
Effective invasion thresholds of CAIN and ClvR drives under gamete viability fitness costs. Baseline germination rate *b* and drive-associated gamete viability cost (*s*_*g*_ ; the probability a drive-carrying gamete is nonviable) were varied, with all other parameters fixed at their default values (Table S2). Blue points represent observed effective invasion thresholds, defined as the minimum introduction frequency resulting in successful drive spread in more than 50% of simulation replicates (see Figures S13 and S14). Red triangles indicate thresholds predicted by 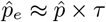. Points at *b* = 1 correspond to baseline invasion thresholds 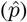 in populations without seedbanks. **a–c**, CAIN effective invasion thresholds for *s*_*g*_ = 0.05, 0.1, 0.25 and *b* = 0.05, 0.2, 0.5, 1. Absence of blue points at these *b* values indicates parameter sets where invasion consistently failed, and absence of red triangles indicates predicted 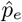 exceeding 1. **d–f**, ClvR effective invasion thresholds, presented as in panels **a**–**c**.

We next explored the effects of a fitness cost reducing seed viability (*s*_*s*_ *>* 0) with *s*_*g*_ = 0. We define *s*_*s*_ as a codominant reduction in the baseline survival rate of seeds (*d*). Thus, the survival rate for a seed of age *a* is *d/a*^*q*^ if wild type, (*d − s*_*s*_*/*2)*/a*^*q*^ if drive heterozygous, and (*d− s*_*s*_)*/a*^*q*^ if drive homozygous, where *q* governs the rate at which survival rates decline with seed age. Importantly, *s*_*s*_ *>* 0 results in genotype-dependent variation in the average seedbank duration (*τ*) and total germination probability (*γ*). As the baseline survival rate decreases, *τ* also decreases because older seeds are more likely to have died, shifting the distribution of germinated seeds toward younger ages. Similarly, the total germination probability *γ* declines with *d* due to reduced survival rates across all seed-age classes. Given these complex genotype-dependent shifts in *τ* and *γ*, we did not expect 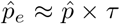 to apply. Nonetheless, we still observed a consistent positive correlation between 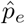 and *τ*, along with drive failure at higher combinations of 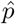 and *τ* (Figures S19–S25). However, we expect the precise relationship among 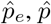, and seedbank parameters to be more complex when drive fitness costs manifest in seeds rather than plants.

## 3 Discussion

In this study, we developed the first comprehensive gene drive modeling framework that explicitly integrates plant-specific life-history traits, including seed dormancy, and investigated the eco-evolutionary dynamics of the CAIN and ClvR drives under realistic ecological conditions. Our results reveal the critical roles of seedbanks in modulating gene drive efficacy, suppression potential, and confinement in plant populations.

In our model, the ClvR male suppression drive consistently outperformed both the ClvR female suppression and CAIN male suppression drives, with the latter two achieving population elimination only under sufficiently low population fecundity or total germination probability (*γ*) (Figures S5, S7, and S10). These results contrast with those of Liu et al. [1] and Oberhofer et al. [2], who reported successful population elimination by ClvR female suppression and CAIN male suppression, respectively, in models without dormancy. We attribute this discrepancy to the lower low-density growth rates (*β*) assumed in their studies compared to ours. Indeed, decreasing *β* to their default values reduces the required genetic load, enabling even weaker constructs (ClvR female suppression and CAIN male suppression) to reach these thresholds and collapse the population. Our model employs relatively high *β* values because we assume complete fertilization of ovules when effective ovules are limiting and pollen is abundant, and complete utilization of pollen when effective pollen is limiting and ovules are abundant. Consequently, when evaluating our model without dormancy, all seeds germinate immediately, and at low densities below carrying capacity, all seedlings develop into plants. However, natural populations often experience seedling mortality due to external biotic and abiotic factors—such as interspecific competition, predation, fungal pathogens, and physical disturbances—independent of seedling density [37–39]. Such factors would decrease *β* below our modeled values, thereby lowering the required genetic load of the drive and potentially allowing weaker drives to achieve population elimination. Thus, our model provides a conservative estimate of suppression drive efficacy, and the required genetic load formula (Eq. 4) may require adjustment to incorporate additional population-specific constraints on seedling establishment and survival.

Several limitations of our model present opportunities for further investigation. We assumed a panmictic population, an annual life cycle, constant seedbank parameters, wind pollination, and the absence of selfing and spatial dynamics. These assumptions underestimate the complexity of natural populations, where genetic variation, environmental fluctuations, and spatial heterogeneity often influence seed survival, germination, pollen dispersal, and drive dynamics. Additionally, we fixed CAIN and ClvR parameters at their experimentally derived values [1, 2]; however, exact drive parameters are likely subject to some degree of uncertainty, especially in species other than *Arabidopsis thaliana*. We also did not model functional resistance alleles that prevent drive cleavage but retain target gene function. Previous theoretical work shows that if functional resistance alleles confer fitness advantages against the drive, they can eventually outcompete the drive allele [2, 40, 41]. However, we expect functional resistance rates to be low for CAIN and ClvR due to their use of multiple guide RNAs (gRNAs), increasing target cleavage redundancy [42–45]. Although resistance alleles may eventually arise in sufficiently large populations, their establishment probability would likely be reduced in plant populations with low germination rates. Another phenomenon known to hinder the success of suppression drives is “chasing”, a recolonization-extinction dynamic in spatially-structured populations in which wild-type individuals repopulate areas previously cleared by the drive, often preventing population elimination [41, 46, 47]. Seedbanks may significantly influence chasing dynamics, especially when coupled with other plant-specific life-history traits, such as long-distance dispersal of pollen and seeds and reduced low-density growth rates of populations with low germination rates. Such interactions could be complex and warrant further investigation.

Our results highlight three key life-history metrics critical to the effective deployment of gene drives in weed populations: (i) average seedbank duration *τ*, (ii) total germination probability *γ*, and (iii) fecundity (mean effective ovule count *n*_*bo*_ and mean effective pollen grain count *n*_*bp*_ per individual). If a candidate population exhibits a long average seedbank duration under natural conditions and controlled experiments without seedbanks indicate a high baseline invasion threshold 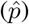, the effective invasion threshold 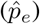 may exceed practical single-release frequencies (e.g., Figures 6c, f and S24c, f). In such cases, multiple consecutive releases at sufficiently high frequencies could potentially enable the drive to overcome seedbank dilution effects, and this extra release effort could improve drive confinement within the target population. For modification drives, our results suggest that fecundity has minimal impact on drive success; however, long seedbank durations can substantially delay drive fixation (Figures 3 and S4). For suppression drives, we found that population elimination is facilitated by reduced fecundity and lower total germination probability, as both decrease the required genetic load of the drive (Figure 4). Low-fecundity populations also benefit from smaller population sizes, thus requiring fewer drive individuals to achieve necessary release frequencies. Lastly, because selfing can inhibit drive spread and evolve as a resistance mechanism [32, 41, 48, 49], we recommend prioritizing dioecious species as deployment targets. However, if the target species is monoecious or hermaphroditic and capable of selfing, we suggest deploying modification drives with minimal fitness costs at higher introduction frequencies to encourage more rapid fixation, thereby minimizing time in which a selfing tendency can evolve.

Although CRISPR-based gene drive systems have made significant advancements in controlling disease-vector mosquitoes [50–54], their potential for agricultural weed control has remained largely unexplored. This gap likely stems from the lack of experimentally validated gene drives in plants, with CAIN and ClvR only recently having been developed [1, 2], and the assumptions that key plant life-history traits (e.g., high fecundity, self-pollination, and seed dormancy) constrain drive efficacy [17, 18]. Our findings challenge this view by demonstrating that seedbanks can, in some cases, enhance both the efficacy and biosafety of gene drives in plant systems. Specifically, seedbanks can lower the required genetic loads for suppression drives and introduce higher invasion thresholds under drive-associated fitness costs, substantially reducing risks of unintended gene drive spillover into non-target populations. Thus, highly efficient gene drive constructs can become inherently safer in populations with seed dormancy. Achieving comparable thresholds in populations without seedbanks would require either complex gene drive architectures, which are often challenging to engineer in practice (e.g., underdominance drives [55–58]), or substantially increased fitness costs of toxin-antidote drives, significantly reducing drive efficacy. Collectively, our work highlights seed dormancy not as a barrier, but as an ecological trait that can be strategically leveraged for gene drive deployment, providing opportunities to optimize feasibility, effectiveness, and safety of genetic interventions in weed populations. More broadly, our study underscores the importance of explicitly integrating ecological and life-history complexity into genetic control strategies to enable safer and more targeted biotechnological solutions.

## 4 Methods

### 4.1 Gene drive systems

We parameterized the CAIN and ClvR gene drives using experimentally derived values from their respective experimental studies [1, 2]. Figure 1 illustrates the mechanism of each drive, and Table S1 summarizes the corresponding model parameters used in our simulations. We modeled the germline cleavage rate as the probability that a wild-type target allele—located on a separate chromosome from the drive—is cleaved in the germline of an individual carrying at least one drive allele. This rate may differ based on the sex of the individual. We assumed the drive carries multiple gRNAs that target conserved sites within the target gene, resulting in an extremely low functional resistance rate [44]; that is, the probability of a target allele becoming uncleavable at every gRNA target site while preserving the gene’s function is minimal. Thus, we do not model functional resistance; all cleavage events are assumed to produce a nonfunctional (disrupted) copy of the gene. We also exclude maternal or paternal Cas9 deposition (i.e., embryo cleavage) and intrinsic drive-associated fitness costs by default, as these were not reported or explored in the referenced studies. However, we do consider drive fitness costs reducing gamete viability and seed survival when evaluating the effect of seedbank parameters on threshold-dependent drives.

The CAIN study [1] reported a germline cleavage rate of 0.984 in males and 0.941 in females. The authors also observed a 96% penetrance rate of the *NPG1* target gene, which we modeled as a 4% probability that a pollen grain lacking a functional copy of the gene remains viable. Because *NPG1* affects only male gametes, ovules with disrupted target alleles are assumed to remain viable. For a modification version of CAIN, we assume that a cargo gene is attached to the drive construct, such that drive carriers have a desirable phenotype. We refer to the modification construct simply as “CAIN”. For suppression, we assume that the CAIN drive is inserted into a haplosufficient essential male fertility gene in a manner that disrupts the gene, such that males with two drive alleles are sterile. We refer to this construct as “CAIN male suppression”.

For ClvR [2], exact estimates for male and female cleavage rates are unavailable. We inferred these values from the male and female drive inheritance rates, averaged across experimental crosses and weighted by sample sizes (Section S1). We estimated a male germline cleavage rate of 0.974. In females, drive inheritance rates were lower than in males, a difference attributed to maternal carryover—the partial deposition of the rescue protein (encoded by the drive) into female gametes. Assuming female cleavage occurs at the same rate as in males, we estimated the maternal carryover rate to be approximately 0.207. We modeled this as a 20.7% probability that a female gamete lacking a functional *YKT61* allele remains viable when produced by a drive-carrying female. For population modification, we assume that a cargo gene is attached to the ClvR drive, such that individuals with a drive allele have some desirable phenotype. We refer to the modification version of the drive as “ClvR”. For population suppression, we consider two variants of ClvR: “ClvR male suppression”, in which the drive is inserted into a haplosufficient, essential male fertility gene in a disruptive manner such that male homozygotes are sterile, and “ClvR female suppression”, in which the drive is inserted into a haplosufficient, essential female fertility gene in a disruptive manner such that female homozygotes are sterile.

### 4.2 Life cycle model of annual plants

Our model is based on wind-pollinated weeds, a major group of potential gene drive targets [17]. While some species can be monoecious or hermaphroditic and self-fertilize to varying degrees, we assume a strictly dioecious system to avoid the potential evolution of selfing as a protective mechanism against gene drive-induced fitness costs [48, 49]. We further assume a panmictic population with an annual life cycle (Figure 2). The model employs a stochastic, agent-based framework that explicitly accounts for the plant life cycle and incorporates the CAIN and ClvR drive systems. Our model extends beyond the deterministic homing drive framework of Barrett et al. [18], allowing for a more biologically and ecologically realistic representation of drive dynamics.

#### Plants to seeds: reproduction (Figure 2a)

In our model, we do not explicitly track all ovules or pollen grains produced in the population; instead, we focus on “effective gametes” that actually contribute to reproduction. We define an “effective ovule” as an ovule capable of fertilization and subsequent development into a viable seed and an “effective pollen” as a pollen grain that can reach a receptive female by wind and successfully germinate. Because the probability of pollen deposition on a receptive stigma is extremely low in wind-pollinated systems [59], the expected effective pollen count is substantially lower than the total pollen production per male. This count varies among males due to differences in pollen production and transfer efficiency [59–61]. Analogously, the effective ovule count varies among females because of resource allocation, developmental anomalies, and other factors affecting ovule viability [62–65]. We account for this individual-level variation by sampling the number of effective gametes per individual from a Poisson distribution with sex-specific means.

We assume that pollen deposition is uniformly distributed among a female’s flowers, such that only the total number of effective ovules per female determines reproductive output. Let *n*_*bo*_ denote the mean effective ovule count per wild-type female, which was held constant throughout the simulation. We draw each fertile female’s maximum number of effective ovules from a Poisson distribution with mean *n*_*bo*_. We then consider drive processes in the germline that created these gametes and whether any such gametes would have been rendered nonviable due to the drive (Section 4.1). For the CAIN drive—whose target gene only affects male gametes—all ovules are viable, so a female’s effective ovule count will always equal her maximum effective ovule count drawn from the Poisson distribution. This is also the case for the ClvR drive when females lack a drive allele and disrupted target allele. However, if a female carries a ClvR drive allele or disrupted target allele, some ovules will be nonviable if they inherit a disrupted target allele with no rescue (through co-inheriting the drive or through maternal carryover), thereby reducing her effective ovule count below her maximum number drawn. For suppression drives, sterile females are assumed to produce no effective ovules and are excluded from reproduction.

We define *n*_*bp*_ as the mean effective pollen count per male in a wild-type population at carrying capacity *K*. The probability that a pollen grain successfully reaches a receptive stigma at time *t* is assumed to be proportional to the number of fertile females *N*_*f*_ (*t*), with all stigmas equally likely to capture pollen. Accordingly, as *N*_*f*_ (*t*) increases, the per-pollen grain probability of successful deposition increases proportionally.

To model this dependency, we scale *n*_*bp*_ by the ratio of *N*_*f*_ (*t*) to the expected number of fertile females at carrying capacity, *K/*2, yielding the time-dependent mean number of effective pollen grains per male, *n*_*p*_(*t*):

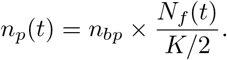

At carrying capacity, *n*_*p*_(*t*) = *n*_*bp*_; however, as the number of fertile females declines, a greater fraction of pollen grains fails to reach receptive females, thereby reducing the effective pollen count. This scaling is especially relevant for suppression drives that reduce the number of fertile females. A similar procedure is conducted in males as in females: for each fertile male, we first sample the maximum effective pollen count from a Poisson distribution with mean *n*_*p*_(*t*), then model drive processes in the germline that produced these pollen grains, and lastly, discard any pollen grains that were rendered nonviable due to the drive. Because of drive-associated loss of gamete viability, males carrying a CAIN or ClvR drive allele or disrupted target allele ultimately produce fewer effective pollen grains than wild-type males. For suppression drives targeting male fertility genes, sterile males are assumed to produce no effective pollen and are excluded from reproduction.

Effective pollen grains are then collected from the *N*_*m*_(*t*) fertile males and randomly allocated to the *N*_*f*_ (*t*) fertile females. The total number of effective pollen 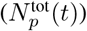, has expectation *N*_*m*_(*t*) *× n*_*p*_(*t*). We model pollen allocation as a multinomial random variable with 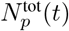 trials and uniform category probabilities 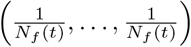, assuming each fertile female is equally likely to receive a pollen grain. The expected number of pollen grains per female is 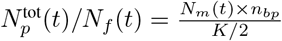, though stochastic variation may lead to deviations from this mean.

Fertile females typically receive enough effective pollen to fertilize all their effective ovules, in which case each effective ovule randomly samples an effective pollen grain from the female’s effective pollen pool, assuming all effective pollen are equally likely to be chosen. However, if a suppression drive reduces pollen availability, some females may not receive enough effective pollen to fertilize all effective ovules. In such case, each effective pollen grain fertilizes a randomly chosen effective ovule from the female, assuming all effective ovules have an equal probability of being fertilized. Fertilized effective ovules become seeds, which have an equal probability of being male or female. Female plants that receive no effective pollen produce no seeds. Similarly, if a suppression drive reduces the number of effective ovules, fewer fertilization events occur, thereby reducing seed production.

#### Movement out of seedbank: age-dependent seed mortality and germination (Figure 2b and c)

Newly produced seeds enter the seedbank and survive their first year with probability *d*. Thereafter, survival rates decline with seed age, such that the probability of an age-*a* seed surviving, given its persistence in the seedbank, is *d/a*^*q*^, where the parameter *q* determines the rate at which survival rates decrease with age. Surviving seeds have an opportunity to germinate each year, with age-dependent germination probabilities modeled similarly to survival probabilities. Newly produced seeds germinate with probability *b*, conditional on surviving their first year. For seeds of age *a >* 1, germination rates decline with age, such that the probability of an age-*a* seed germinating, given its persistence in the seedbank, is *b/a*^*m*^, where *m* determines the rate at which germination rates decrease with age. Germination rates are assumed to be genotype-independent. Seeds can persist in the seedbank for a maximum of *L* years [66, 67], after which they are assumed to be nonviable and are removed from the model. To simulate populations without seedbanks, we set *d* = 1 and *b* = 1, such that all seeds survive and immediately germinate. In such a case, seed age never exceeds one year, making parameters *m, q*, and *L* have no effect.

#### Seeds to plants: seedling competition (Figure 2d)

Germinated seeds develop into seedlings that compete for limited resources [39, 68–70]. We assume that seedling competition regulates the population size, maintaining the number of plants near the carrying capacity *K* at equilibrium. We denote the number of seedlings at year *t* with *N*_sdl_(*t*). When *N*_sdl_(*t*) *≤ K*, all seedlings survive to adulthood. When *N*_sdl_(*t*) *> K*, density-dependent competition reduces survival probability proportionally. The probability of a seedling surviving to adulthood at time *t* is given by:

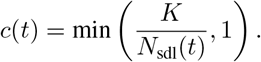

Surviving seedlings mature into adult plants, completing the life cycle. Each model time step corresponds to one year.

### 4.3 Average seedbank duration and total germination probability

To obtain the average seedbank duration (*τ*), we first derive the probability of germinating at age *a* (*g*_*a*_) for seeds of age *a ∈ {* 1, …, *L*}, following previous approaches [30, 35, 71]. For a seed to germinate at age *a*, it must (i) remain dormant through the preceding *a −* 1 years without dying or germinating and (ii) both survive and germinate in year *a*. Thus, *g*_*a*_ is given by

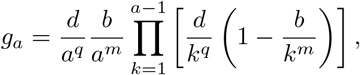

where *d* and *b* are the baseline survival and germination rates, and *q* and *m* are shape parameters governing the age-dependent decline in survival and germination, respectively, as defined in Section 4.2.

We define the total germination probability *γ* as the cumulative probability of germination across all seed ages up to the maximum seed age *L*:

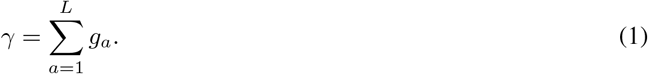

With this, we derive the average seedbank duration *τ*, equivalent to both the expected age of germinated seeds and the generation time of the population:

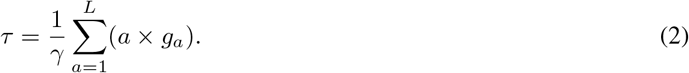

### 4.4 Implementation of gene drive simulation model with a seedbank

We implemented our model in SLiM v4.0.1 [34], an individual-based, forward-time population genetic simulation framework. We initialized each simulation with *K* wild-type plants and allowed the plant and seed populations to equilibrate for 2*L* years before introducing the drive. This burn-in period was sufficient for the empirical average seedbank duration to match its analytical expectation (Section 4.3). We then introduced the drive at a frequency of *p*_0_ by randomly converting a fraction 2*p*_0_ of plants into drive heterozygotes. The drive was only introduced once per simulation. The simulation was run until one of the following outcomes occurred: drive fixation, drive loss, population suppression, or 500 years elapsed after drive introduction.

### 4.5 Low-density growth rate

For modification drives, we evaluated the probability of fixation and time to fixation. For suppression drives, we evaluated the probability and timing of population elimination. In the latter case, the efficacy of suppression depends largely on the density-dependent dynamics of the population [72, 73]. As the suppression drive spreads, gamete production declines, but the few remaining gametes face reduced competition and an increased probability of successful fertilization. If the maximum per-capita growth rate at low density, denoted *β*, is sufficiently high, population elimination may be prevented. To derive *β* for our suppression constructs, we first note that maintaining the population at carrying capacity *K* requires, on average, each female to produce at least two fertilized effective ovules (assuming abundant effective pollen grains) or each male to pollinate two effective ovules (assuming abundant effective ovules). These fertilized ovules become seeds, which then must survive the seedbank and germinate. Therefore, the minimum required seed production per male or female is *n*_min_ = 2*/γ*, where *γ* is the total germination probability, as defined in Eq. 1.

For suppression drives that generate sterile males, the male low-density growth rate *β*_*m*_ can be derived by considering the limiting case in which a single fertile male remains, while the number of fertile females remains at *K/*2. This male produces, on average, *n*_*bp*_ viable pollen grains. Given an excess of effective ovules and no competition from other males, all effective pollen is assumed to fertilize effective ovules successfully, producing *n*_*bp*_ seeds per male at low density. Thus, the male low-density growth rate is *β*_*m*_ = *n*_*bp*_*/n*_min_ = *n*_*bp*_*γ/*2. Alternatively, if the suppression drive generates sterile females, the female low-density growth rate *β*_*f*_ is derived by considering the limiting case in which a single fertile female remains, while the number of fertile males is *K/*2. This female produces *n*_*bo*_ effective ovules on average. With abundant effective pollen and no competition among females, all of her effective ovules are successfully fertilized, resulting in *n*_*bo*_ seeds produced per female. The corresponding female low-density growth rate is *β*_*f*_ = *n*_*bo*_*/n*_min_ = *n*_*bo*_*γ/*2.

### 4.6 Required genetic load

We focus on the low-density growth rate (*β*) that the suppression drive must overcome to achieve suppression. Let *n* denote the baseline number of effective gametes per wild-type individual. Because the parameters determining reproductive success differ according to which sex is rendered sterile by the drive, we set these parameters sex-specifically: for drives inducing sterile males, we set *β* = *β*_*m*_ and *n* = *n*_*bp*_, and for drives inducing sterile females, we set *β* = *β*_*f*_ and *n* = *n*_*bo*_. The suppressive effect of the drive can be quantified by its genetic load, which measures the reduction in population fitness induced by the drive [72, 74–77]. We measure population fitness by the total number of effective gametes produced in a given year, relative to the number expected in a wild-type population of the same size. We define the genetic load *λ*(*t*) at time *t* as:

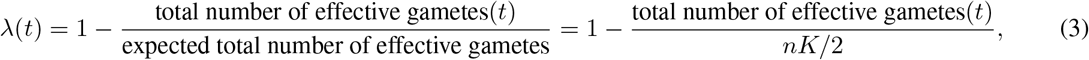

where *nK/*2 represents the expected number of effective gametes, assuming the population is at carrying capacity.

For the drive to suppress the population, it must reduce the effective gamete pool below the threshold necessary to produce *n*_min_ seeds per individual. The critical genetic load required for population decline is at least 1 *−* 1*/β* [75], which we define as the *required genetic load λ*^*\**^:

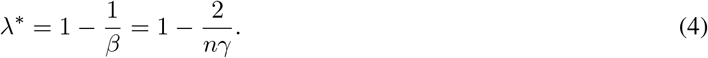

### 4.7 Fitness cost of gene drive

Our base model assumes no fitness cost associated with CAIN or ClvR. To investigate the impact of drive-associated fitness costs, we extended the model to incorporate drive-induced fitness costs on gamete viability and seed survival. We define *s*_*g*_ as the probability that a drive-carrying gamete is nonviable, effectively representing a codominant fitness cost reducing gamete production. Thus, *s*_*g*_*/*2 of drive-heterozygotes gametes and *s*_*g*_ of drive-homozygotes gametes are nonviable in expectation. Similarly, we define *s*_*s*_ as a codominant fitness cost reducing seed survival, modeled as a reduction in the baseline survival rate (*d*). Drive-heterozygous seeds have a baseline survival rate of *d − s*_*s*_*/*2, while drive-homozygous seeds have *d − s*_*s*_. Seed survival rates also decline with seed age (*a*), following the general form: *d/a*^*q*^ for wild-type, (*d − s*_*s*_*/*2)*/a*^*q*^ for drive heterozygotes, and (*d − s*_*s*_)*/a*^*q*^ for drive homozygotes, where *q* controls the rate at which survival probabilities decline with age. We examined each fitness cost separately to quantify its impact on the effective invasion threshold of the drive.

## Data and Code Availability

The data and code used for the simulation studies presented in this manuscript is publicly available on Zenodo at doi:10.5281/zenodo.15110717.

## Acknowledgments

IKK, BCH, and PWM were supported by the National Institutes of Health under award R35GM152242. RC was supported by the National Science Foundation Graduate Research Fellowship under award DGE2139899. JC was supported by the Center for Life Sciences and by the National Natural Science Foundation of China grants 32270672 and W2432018. JK was supported by the National Institute of General Medical Sciences of the National Institutes of Health under award R35GM156957.

