## Supplementary Information for "Gene drive dynamics in plants: the role of seedbanks"

### SI Text

#### S1 Drive parameterization

We obtained the male germline cleavage rate, female germline cleavage rate, and penetrance rate of CAIN directly from Liu et al. [1]. Since these rates were not explicitly reported for the ClvR drive, we inferred them from the experimental data, summary statistics, and interpretations provided by the authors.

We focused on the ClvR-ubq7 construct as this was the most efficient construct tested. We first estimated the male ($p_{d,m}$) and female ($p_{d,f}$) ClvR drive inheritance rates by calculating the sample size-weighted averages from crosses reported in Figures 2 and 3 of Oberhofer et al. [2]. Specifically, $p_{d,m}$ was derived from crosses of drive-heterozygous males with wild-type females, and $p_{d,f}$ from crosses of drive-heterozygous females with wild-type males.

The drive inheritance rates through males ($p_{d,m}$) and females ($p_{d,f}$) are calculated as:

$$\begin{aligned} p_{d,m} &= \frac{\text{number of seeds with drive allele}}{\text{total number of seeds}}, \\ &= \frac{1 \times 357 + 0.988 \times 482 + 0.988 \times 497 + 0.971 \times 513 + 0.993 \times 583 + 0.992 \times 488 + 0.984 \times 685}{357 + 482 + 497 + 513 + 583 + 488 + 685}, \\ &= 0.987. \end{aligned}$$

$$\begin{aligned} p_{d,f} &= \frac{\text{number of seeds with drive allele}}{\text{total number of seeds}}, \\ &= \frac{0.798 \times 257 + 0.858 \times 302 + 0.920 \times 250 + 0.927 \times 303 + 0.789 \times 375 + 0.773 \times 335 + 0.740 \times 407}{257 + 302 + 250 + 303 + 375 + 335 + 407}, \\ &= 0.821. \end{aligned}$$

We assume a high germline cleavage rate, such that the drive heterozygotes involved in Figures 2 and 3 of Oberhofer et al. [2] inherited a disrupted target allele from their drive-carrying parent. Thus, these individuals are heterozygous at the target locus, carrying one disrupted and one intact (non-disrupted) allele.

The authors attributed failed drive inheritance through males to a lack of germline cleavage. They also assumed full penetrance of the *YKT61* target gene, such that pollen carrying a disrupted target allele but no drive allele are always nonviable. Under these assumptions, we can derive the male germline cleavage rate ($c_m$). A drive-heterozygous male produces gametes of which half inherit the drive allele, remaining viable. The other half inherit the wild-type allele, with equal probabilities of inheriting either the disrupted or intact target allele. The non-disrupted target allele is cleaved with probability $c_m$, and cleavage is assumed to always produce a disrupted allele, resulting in gamete nonviability. Thus, the expected male drive inheritance rate ($p_{d,m}$) is given by:

$$\begin{aligned} p_{d,m} &= \frac{\text{fraction of gametes with the drive}}{\text{fraction of viable gametes}}, \\ &= \frac{\frac{1}{2}}{\frac{1}{2} + \frac{1}{2} \times \frac{1}{2} \times (1 - c_m)}, \\ &= \frac{2}{3 - c_m}. \end{aligned}$$

Using our estimated $p_{d,m} = 0.987$, we solve for $c_m$ to obtain a male germline cleavage rate of 0.974.

The authors attributed the lower drive inheritance rates observed through females to maternal carryover rescue: drive-heterozygous females may deposit sufficient rescue protein into their ovules so that an ovule carrying a disrupted target allele but no drive allele remains viable. We denote the maternal carryover rate by $m_r$ and the female germline cleavage rate by $c_f$. A drive-heterozygous female produces gametes as follows: half inherit the drive allele and are always viable, while the other half inherit a wild-type allele. Among the remaining wild-type gametes, half inherit the disrupted target allele, of which a fraction $m_r$ remain viable. The other half inherit a non-disrupted target allele, subject to germline cleavage at rate $c_f$, which, if cleaved, are viable at rate $m_r$. Thus, the expected female drive inheritance rate ($p_{d,f}$)

is:

$$
\begin{aligned}
p_{d,f} &= \frac{\text{fraction of gametes with the drive}}{\text{fraction of viable gametes}}, \\
&= \frac{\frac{1}{2}}{\frac{1}{2} + \frac{1}{2} \times \frac{1}{2} \times m_r + \frac{1}{2} \times \frac{1}{2} \times (1 - c_f) + \frac{1}{2} \times \frac{1}{2} \times c_f \times m_r}, \\
&= \frac{2}{3 - c_f + m_r(1 + c_f)}.
\end{aligned}
$$

Assuming equal germline cleavage rates ($c_f = c_m = 0.974$), we use our observed female drive inheritance rate ($p_{d,f} = 0.821$) to solve for $m_r$, obtaining a maternal carryover rate of 0.207.

### SI Tables

**Table S1. Parameters for the CAIN and ClvR gene drive.** Cleavage, penetrance, and maternal carryover rates were fixed at values estimated from experimental studies [1, 2] and not varied in simulations. Unless stated otherwise, simulations assume no fitness costs ($s_g = s_s = 0$) and an initial drive frequency $p_0$ of 0.1. Fitness costs reducing gamete viability ($s_g$) or seed survival ($s_s$) were systematically varied and examined separately for threshold-dependent drives, as they affect drive introduction thresholds. To assess the invasion threshold under these fitness costs ($s_g > 0$ or $s_s > 0$), we explored the introduction frequency $p_0$. The baseline invasion threshold ($\hat{p}$) is defined as the minimum introduction frequency required for successful drive invasion in populations without a seedbank, and the effective invasion threshold ($\hat{p}_e$) as the analogous threshold in populations with a seedbank. Entries indicated by "–" represent variables without default values because they were either systematically varied or estimated directly from simulations.

| Description | Default Value | Note |
|---|---|---|
| **CAIN [1]** | | |
| Male germline cleavage rate | 0.984 | Fixed |
| Female germline cleavage rate | 0.941 | |
| Target gene penetrance rate | 0.960 | |
| Maternal carryover rate | 0 | |
| **ClvR [2]** | | |
| Male germline cleavage rate | 0.974 | Fixed |
| Female germline cleavage rate | 0.974 | |
| Target gene penetrance rate | 1 | |
| Maternal carryover rate | 0.207 | |
| **Threshold-dependent drives** | | |
| Drive fitness cost reducing gamete viability ($s_g$) | – | Varied across $\{0, 0.05, 0.1, 0.25\}$ |
| Drive fitness cost reducing seed survival ($s_s$) | – | Varied across $\{0, 0.05, 0.1, 0.25\}$ |
| Introduction frequency ($p_0$) | – | Varied over $[0, 1]$ in 0.01 steps (modification drive)<br>Varied over $[0, 0.5]$ in 0.01 steps (suppression drive) |
| Baseline invasion threshold ($\hat{p}$)<br>Effective invasion threshold ($\hat{p}_e$) | – | Estimated directly from simulations |

**Table S2. Parameters, time-dependent state variables, and derived population metrics in the plant life-cycle model.** Default parameter values are listed along with the ranges explored in simulations. Seedbank parameters characterize seed survival and germination dynamics; demographic parameters define population reproductive potential under two fecundity scenarios (the default fecundity setting and the low fecundity setting). Time-dependent variables track the dynamic state of the population composition and drive-induced genetic load throughout simulations. Population metrics were derived analytically from the seedbank and demographic parameters and describe key ecological and genetic features of the modeled population. Entries marked with "–" indicate variables calculated dynamically or parameters without default values due to systematic variation. Note that $m$ was systematically varied in heatmaps but was set to 0 for trajectory plots exploring only $b$.

| | Description | Default Value | Note |
|---|---|---|---|
| | **Seedbank parameters** | | |
| $L$ | Maximum seed age | 10 | Fixed |
| $b$ | Baseline germination rate | – | Varied over [0.05, 1] in 0.05 steps |
| $m$ | Age-dependence germination parameter | 0 | Varied over [0, 2] in 0.1 steps |
| $d$ | Baseline survival rate | 1.0 | Fixed |
| $q$ | Age-dependence survival parameter | 0.1 | Fixed |
| | **Demographic parameters** | | |
| $K$ | Carrying capacity | 10000 | Fixed |
| $n_{bo}$ | Mean effective ovule count per female | 60 | 20 for low fecundity setting |
| $n_{bp}$ | Mean effective pollen count per male at wild-type equilibrium | 180 | 60 for low fecundity setting |
| | **Time-dependent variables** | | |
| $N_{\text{sdl}}(t)$ | Number of seedlings at time $t$ | – | |
| $N_f(t)$ | Number of fertile female plants at time $t$ | | |
| $N_m(t)$ | Number of fertile male plants at time $t$ | | |
| $N_p^{\text{tot}}(t)$ | Total number of effective pollen grains at time $t$ | | |
| $n_p(t)$ | Mean number of effective pollen grains per male at time $t$ | | $n_p(t) = n_{bp} \times \frac{N_f(t)}{K/2}$ |
| $c(t)$ | Seedling survival rate at time $t$ | | $c(t) = \min\left(\frac{K}{N_{\text{sdl}}(t)}, 1\right)$ |
| $\lambda(t)$ | Genetic load imposed by the drive at time $t$ | | Eq. 3 |
| | **Population metrics** | | |
| $g_a$ | Probability of germinating at age $a$ | – | $g_a = \frac{d}{a^q}\frac{b}{a^m}\prod_{k=1}^{a-1}\left[\frac{d}{k^q}\left(1-\frac{b}{k^m}\right)\right]$ |
| $\gamma$ | Total germination probability | | Eq. 1 |
| $\tau$ | Average seedbank duration | | Eq. 2 |
| $n_{\min}$ | Minimum number of seeds per male or female to maintain carrying capacity | | $n_{\min} = 2/\gamma$ |
| $\beta$ | Low-density growth rate of the population | | $\beta_m = n_{bp}\gamma/2$ (male); $\beta_f = n_{bo}\gamma/2$ (female) |
| $\lambda^*$ | Required genetic load to eliminate the population | | Eq. 4 |

#### SI Figures

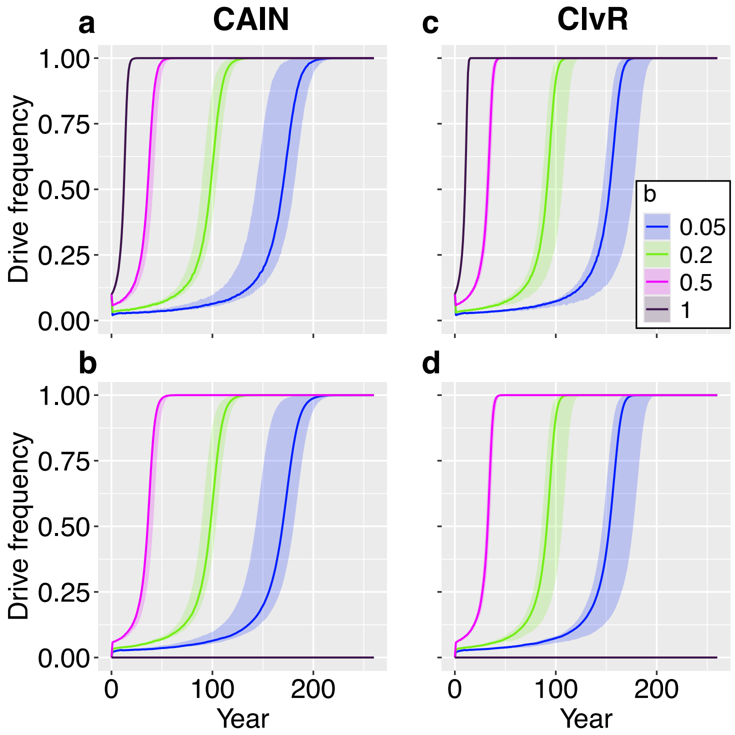


**Figure S1. Frequency trajectories of CAIN and ClvR modification drives in plants and seeds. a,** Frequency trajectories of the CAIN drive in plants under four baseline germination rates ($b$), with other parameters set to default values (Table S2). Solid lines represent median trajectories, and shaded regions indicate the observed range (minimum–maximum) across 10 replicates. **b,** Same as **a** but in seeds. **c–d,** Same as **a–b** but for ClvR drives.

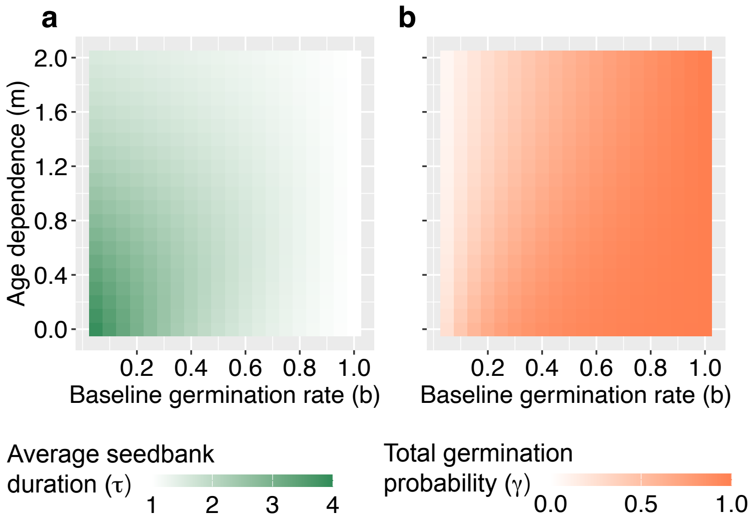


**Figure S2. Average seedbank duration and total germination probability. a,** Average seedbank duration ($\tau$; Eq.2) across varying baseline germination rates ($b$) and age-dependent germination parameters ($m$). All other parameters were set to their default values (Table S2). **b,** Total germination probability ($\gamma$; Eq.1) across varying baseline germination rates ($b$) and age-dependent germination parameters ($m$). All other parameters were set to default values (Table S2).

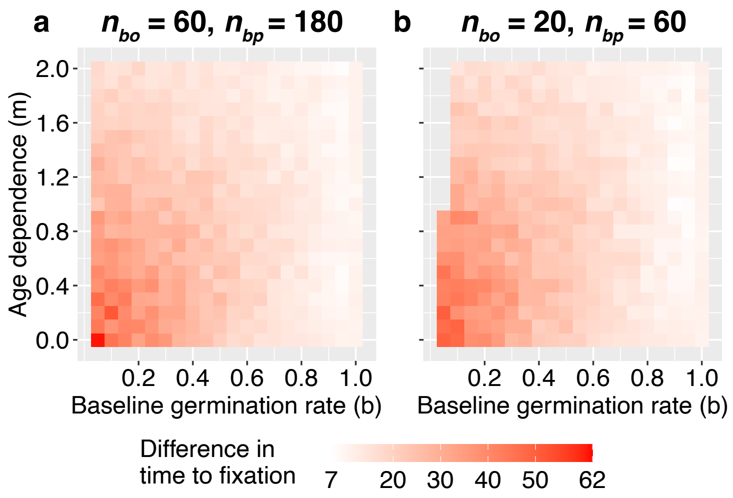


**Figure S3. The difference in fixation time for CAIN and ClvR modification drives. a,** Difference between the average fixation time of CAIN and that of ClvR, calculated as CAIN minus ClvR, under varying baseline germination rates ($b$) and age-dependent germination parameters ($m$). All other parameters were set to their default values in Table S2. Fixation is defined as reaching a 100% drive frequency in both plants and seeds. **b,** Same as **a** but under reduced fecundity conditions, where wild-type females at equilibrium produce an average of 20 effective ovules and wild-type males produce an average of 60 effective pollen grains. Results in the upper-left region of **b** are omitted because low total germination probability ($\gamma$) and seed production lead to an insufficient number of germinated seeds within this parameter space, resulting in population collapse. For fixation times of CAIN and ClvR across this parameter range, see Figures S4.

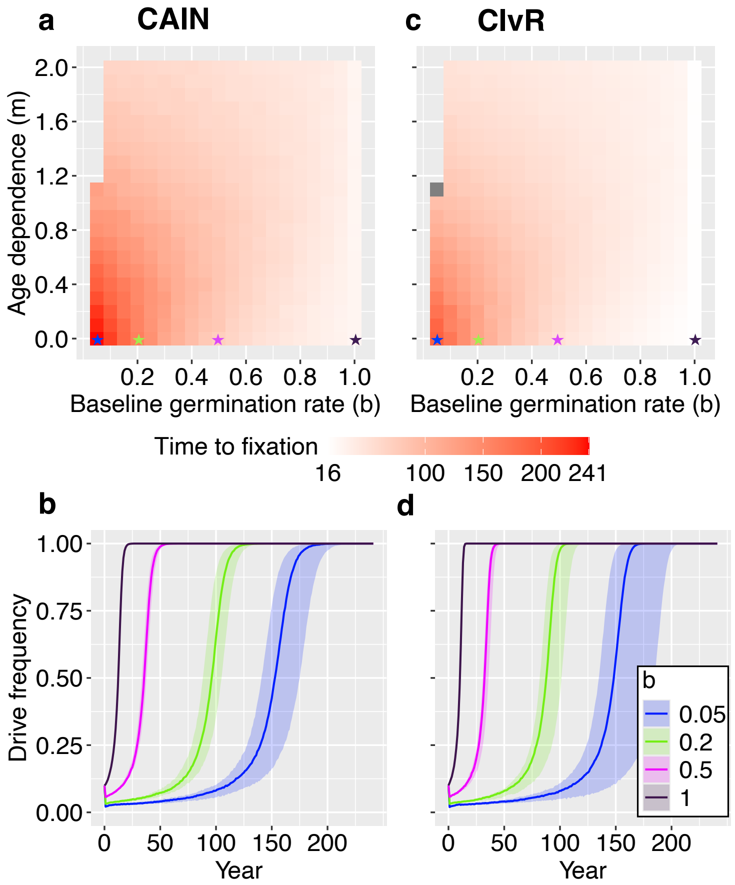


**Figure S4. Spread of CAIN and ClvR modification drives under varying seedbank parameters with reduced fecundity.** Figure design follows Figure 3, except with reduced fecundity: at wild-type equilibrium, females produce an average of 20 effective ovules ($n_{bo}$) and males produce an average of 60 effective pollen grains ($n_{bp}$). Results in the upper-left regions of panels **a** and **c** are omitted because low total germination probabilities ($\gamma$) combined with reduced seed production lead to an insufficient number of germinated seeds, resulting in population collapse before drive introduction.

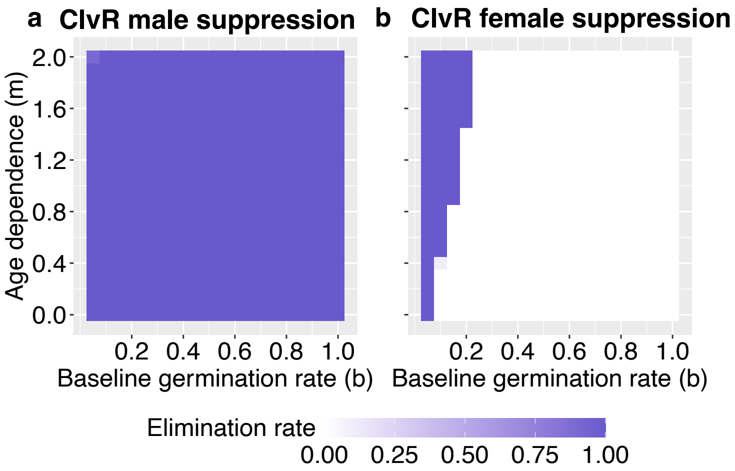


**Figure S5. Population elimination rate of ClvR suppression drives. a,** Fraction of replicates (out of 10) in which the ClvR male suppression drive successfully eliminated the population, across varying baseline germination rates ($b$) and age-dependence germination parameters ($m$). All other parameters were kept at their default values (Table S2). **b,** Same as **a** but for the ClvR female suppression drive.

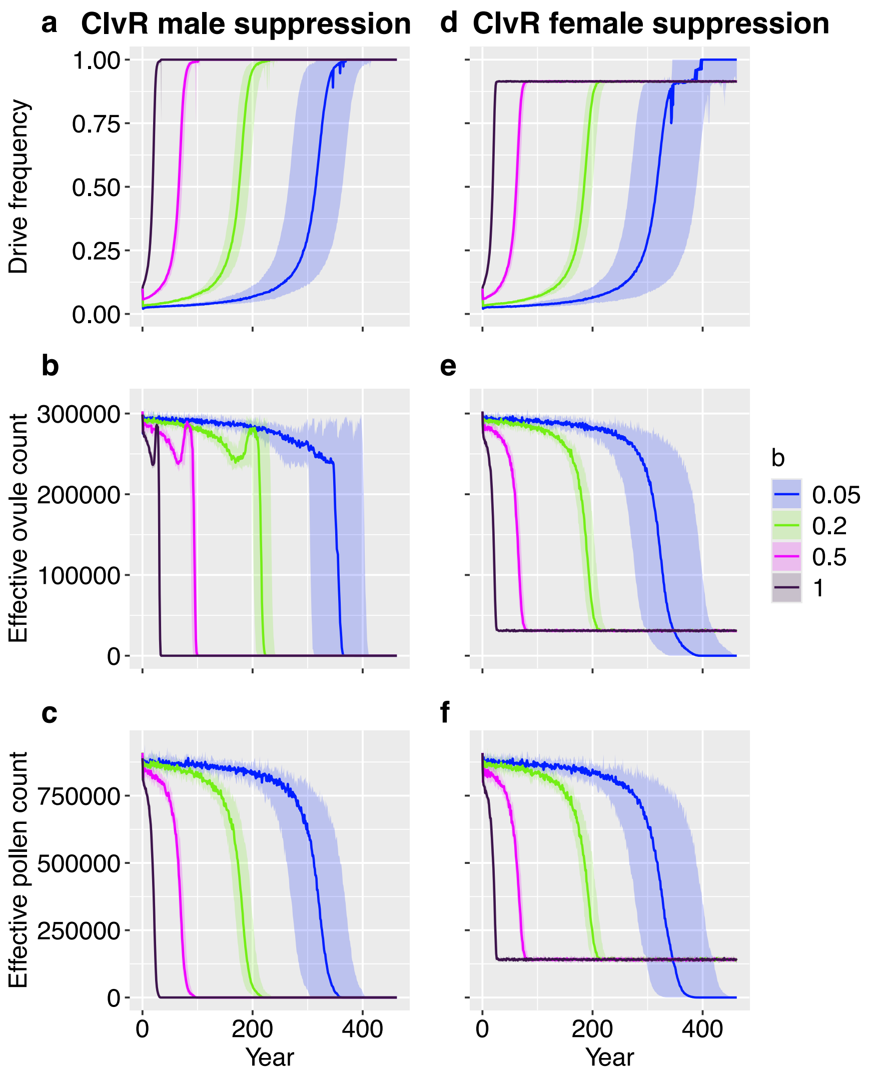


**Figure S6. Trajectories of drive frequency and effective gamete counts for ClvR suppression drives. a,** Trajectory of the ClvR male suppression drive frequency in plants under four different baseline germination rates ($b$), with all other parameters set to their default values (Table S2). Solid lines represent median trajectories, and shaded regions correspond to the observed range (minimum–maximum) across 10 replicates. **b,** Same as **a** but showing the total number of effective ovules in the population. **c,** Same as **a** but showing the total number of effective pollen grains in the population. **d–f,** Same as **a–c** but for the ClvR female suppression drive.

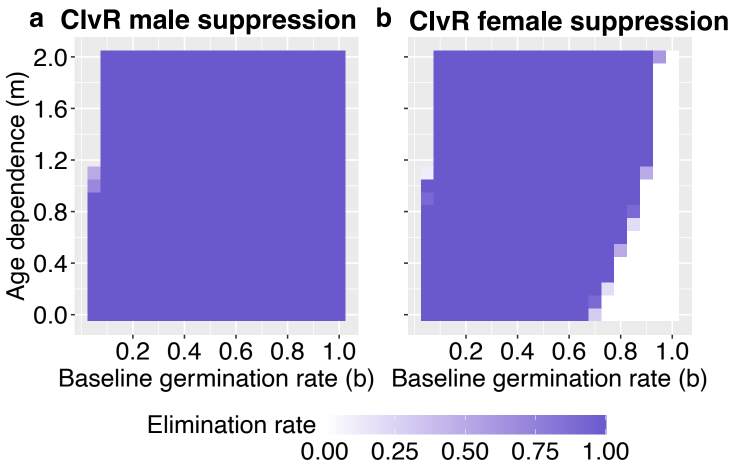


**Figure S7. Population elimination rate of ClvR suppression drives under reduced fecundity. a,** ClvR male suppression. **b,** ClvR female suppression. Figure design follows Figure S5 but with reduced population fecundity: females produce an average of 20 effective ovules, and males produce an average of 60 effective pollen grains at wild-type equilibrium. Results in the upper-left regions of the heatmaps are omitted because, within this parameter space, total germination probability ($\gamma$) and seed production are insufficient to maintain the population.

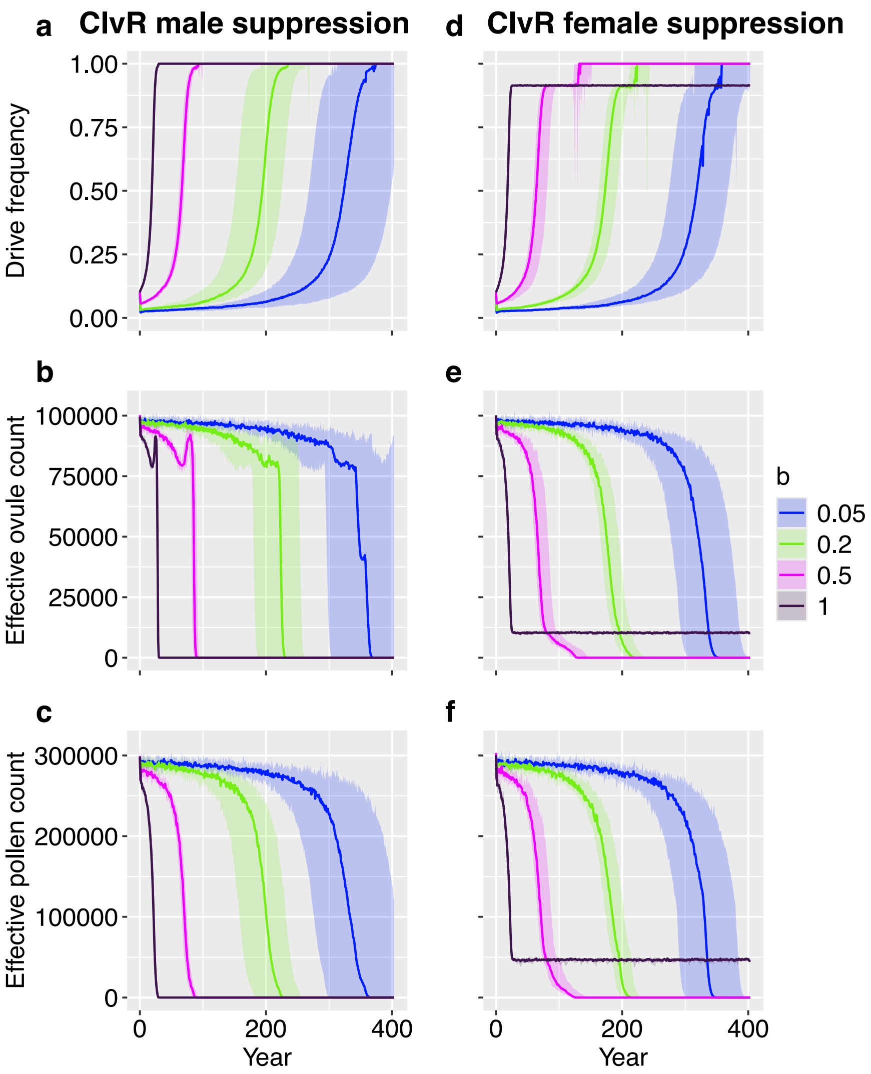


**Figure S8. Trajectories of ClvR suppression drive frequency and effective gamete counts under reduced population fecundity.** Figure design follows Figure S6 but with a reduced mean number of average effective ovule count per female ($n_{bo} = 20$) and a reduced mean number of average effective pollen count per male at wild-type equilibrium ($n_{bp} = 60$).

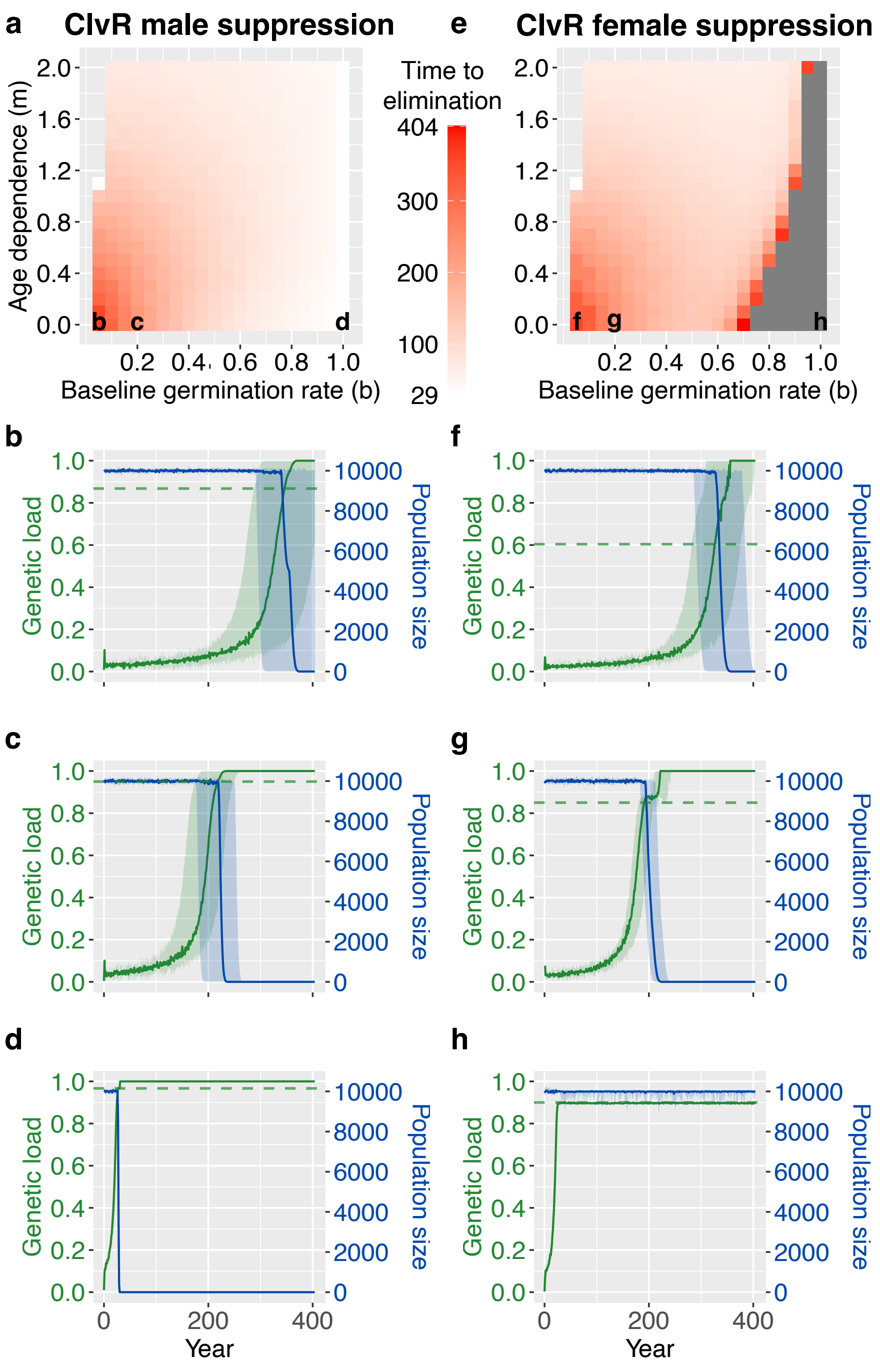


**Figure S9. Dynamics of ClvR suppression drives under varying seedbank parameters and with reduced population fecundity.** Figure design follows Figure 5, except population fecundity was reduced so that wild-type females at equilibrium produce an average of 20 effective ovules, and males produce an average of 60 effective pollen grains. Results in the upper-left regions of panels **a** and **e** are omitted because the combination of low total germination probability ($\gamma$) and low seed production in this range results in an insufficient number of germinated seeds, leading to population collapse before drive introduction.

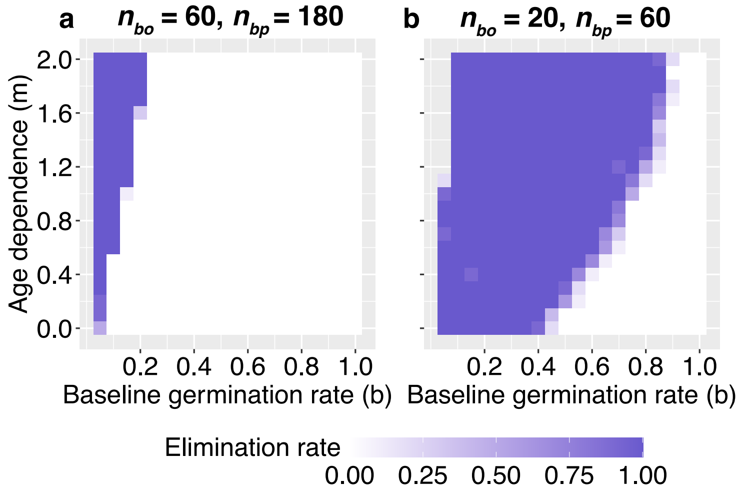


**Figure S10. Population elimination rate of CAIN male suppression under different population fecundity regimes.** Figure design follows Figure S5. **a,** Population elimination rates under CAIN male suppression at default fecundity, where wild-types individuals produce an average of 60 effective ovules per female and 180 effective pollen grains per male at equilibrium. **b,** Same as **a**, with wild-type equilibrium averages of 20 effective ovules per female and 60 effective pollen grains per male. Results in the upper-left region of **b** are omitted because low total germination probability ($\gamma$) combined with reduced seed production yields insufficient germinated seeds, resulting in population collapse before drive introduction.

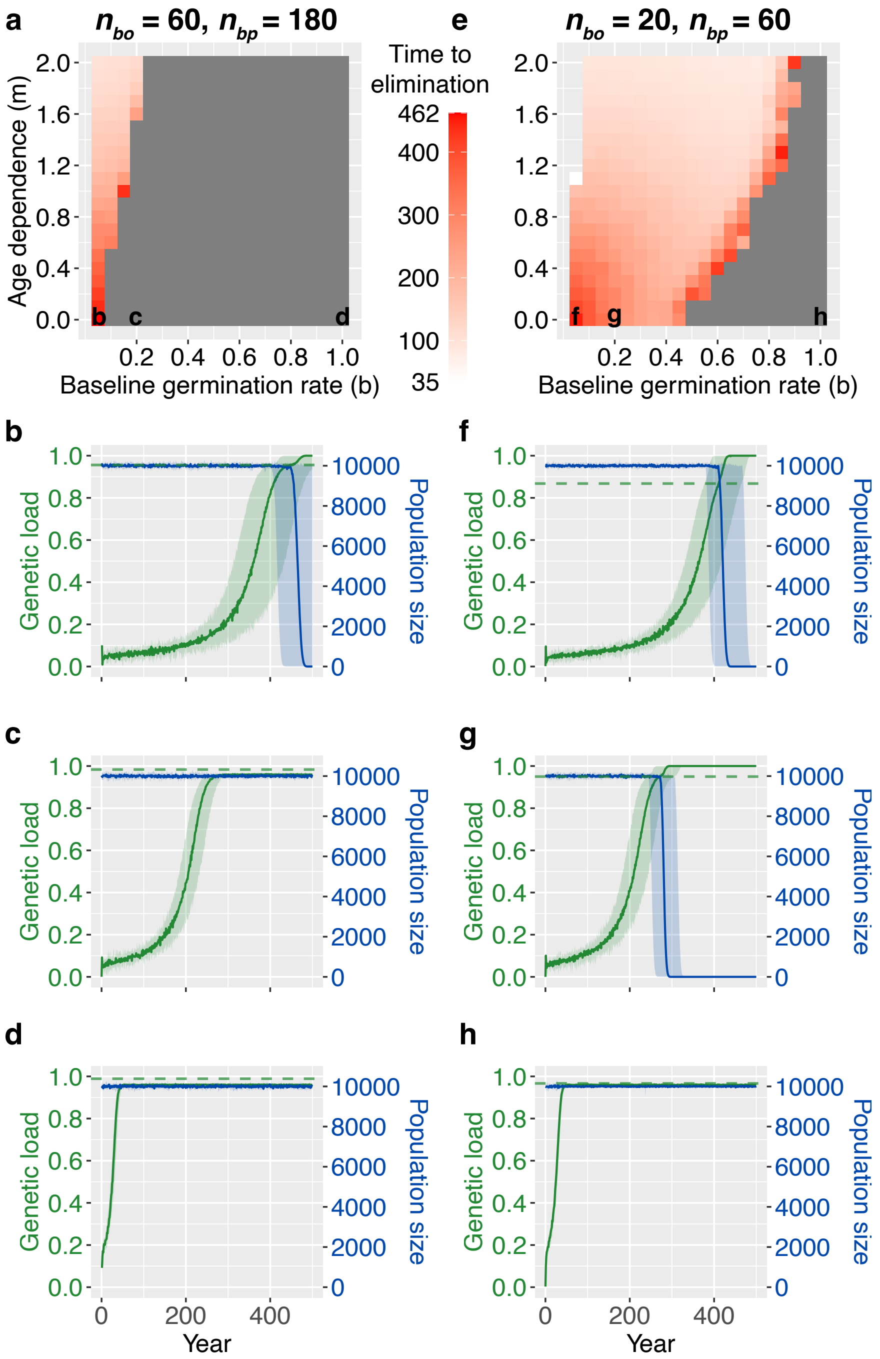


**Figure S11. Dynamics of CAIN male suppression drives under varying seedbank parameters and population fecundity regimes.** Figure design follows Figure 5. The left column shows results under default fecundity, where wild-types individuals produce an average of 60 effective ovules per female and 180 effective pollen grains per male at equilibrium; the right column shows results under reduced fecundity, with average equilibrium production of 20 effective ovules per female and 60 effective pollen grains per male. Results in the upper-left regions of **a** and **e** are omitted because low total germination probability ($\gamma$) combined with reduced seed production yields insufficient germinated seeds, resulting in population collapse before drive introduction.

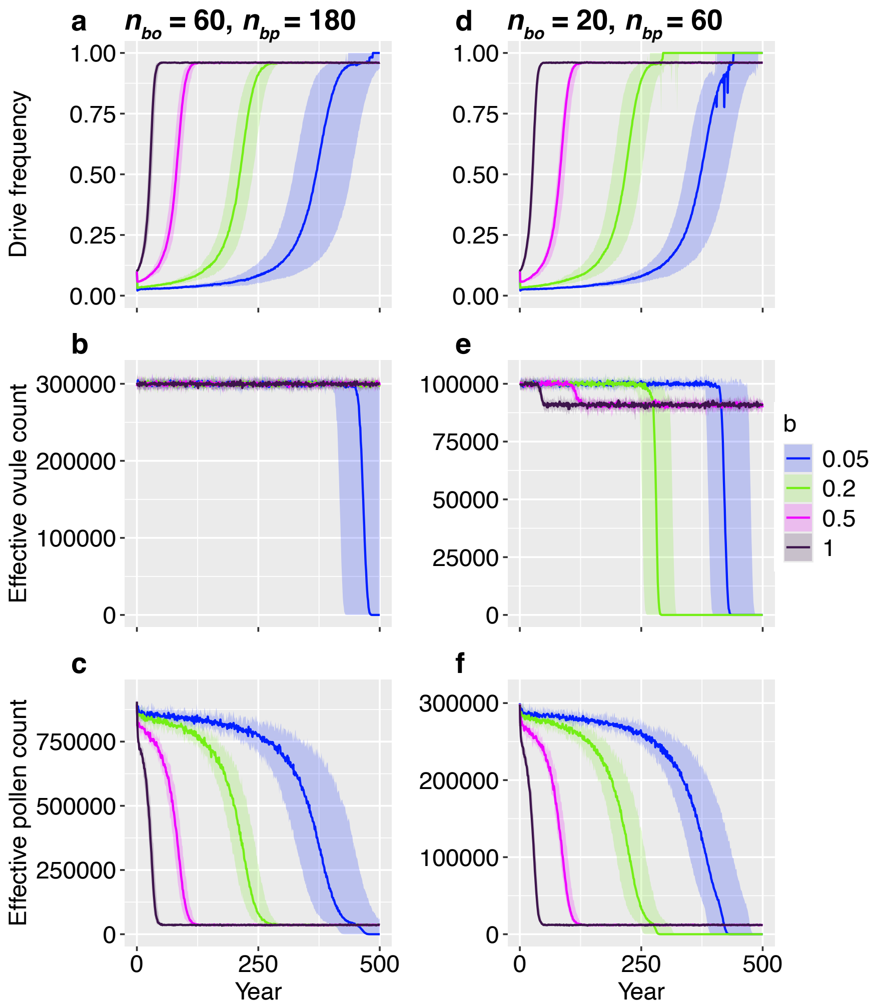


**Figure S12. Trajectories of drive frequency and effective gamete counts for CAIN male suppression under different fecundity regimes.** Figure design follows Figure S6. Panels **a–c** show results under default fecundity, with wild-type equilibrium production averaging 60 effective ovules per female and 180 effective pollen grains per male. Panels **d–f** show results under reduced fecundity, with wild-type equilibrium averages of 20 effective ovules per female and 60 effective pollen grains per male.

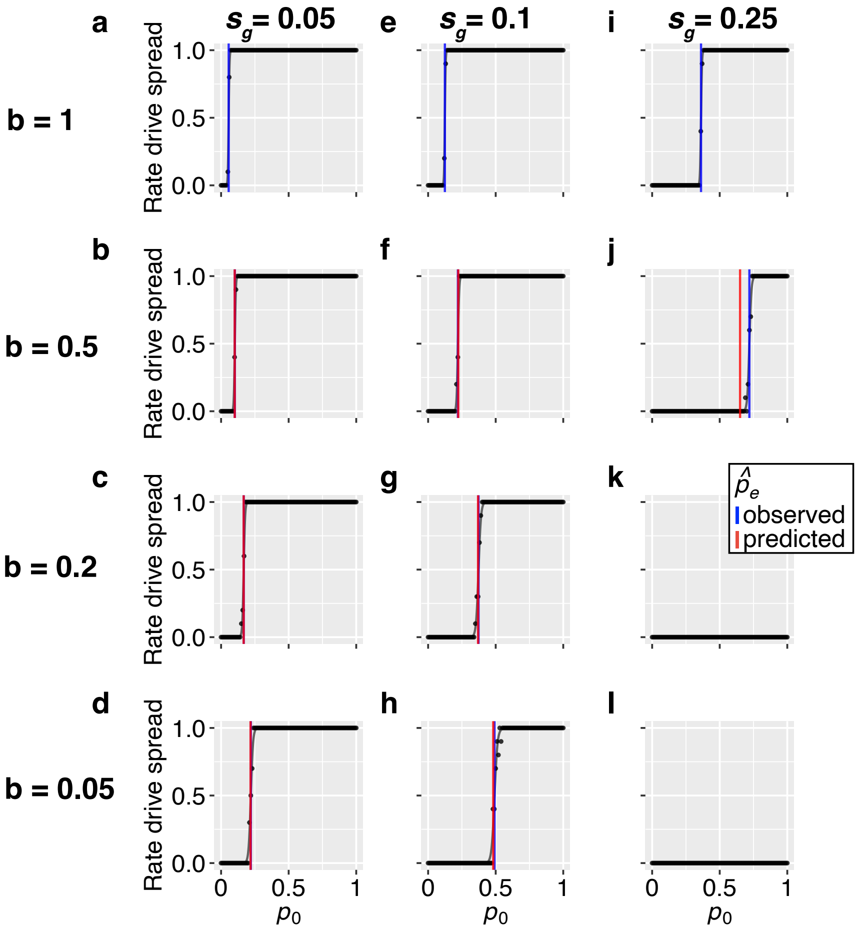


**Figure S13. Invasion ability of the CAIN modification drive under varying gamete viability fitness costs and baseline germination rates.** For a given baseline germination rate ($b$) and drive fitness cost reducing gamete viability ($s_g$, defined as the probability that a drive-carrying gamete is nonviable), the drive was introduced by converting a fraction $p_0$ of plants into drive homozygotes. All other parameters were kept at their default values (Table S2). Each point represents the fraction of replicates (out of 10 total) in which CAIN spread successfully from a given introduction frequency ($p_0$). Drive spread is defined as reaching a final drive frequency that exceeds $p_0$. Logistic curves (blue) were fit to replicate outcomes using the `drc` R package [3], with the effective invasion threshold (blue vertical line) corresponding to the introduction frequency at which CAIN spreads in 50% of replicates (i.e., where the logistic curve intersects with $y = 0.5$). Predicted effective invasion thresholds (red vertical line) were calculated as $\hat{p}_e \approx \hat{p} \times \tau$; values exceeding 1 are not shown. **a–d,** $s_g = 0.05$ and $b = 0.05, 0.2, 0.5, 1$. **e–h,** $s_g = 0.1$ and $b = 0.05, 0.2, 0.5, 1$. **i–l,** $s_g = 0.25$ and $b = 0.05, 0.2, 0.5, 1$.

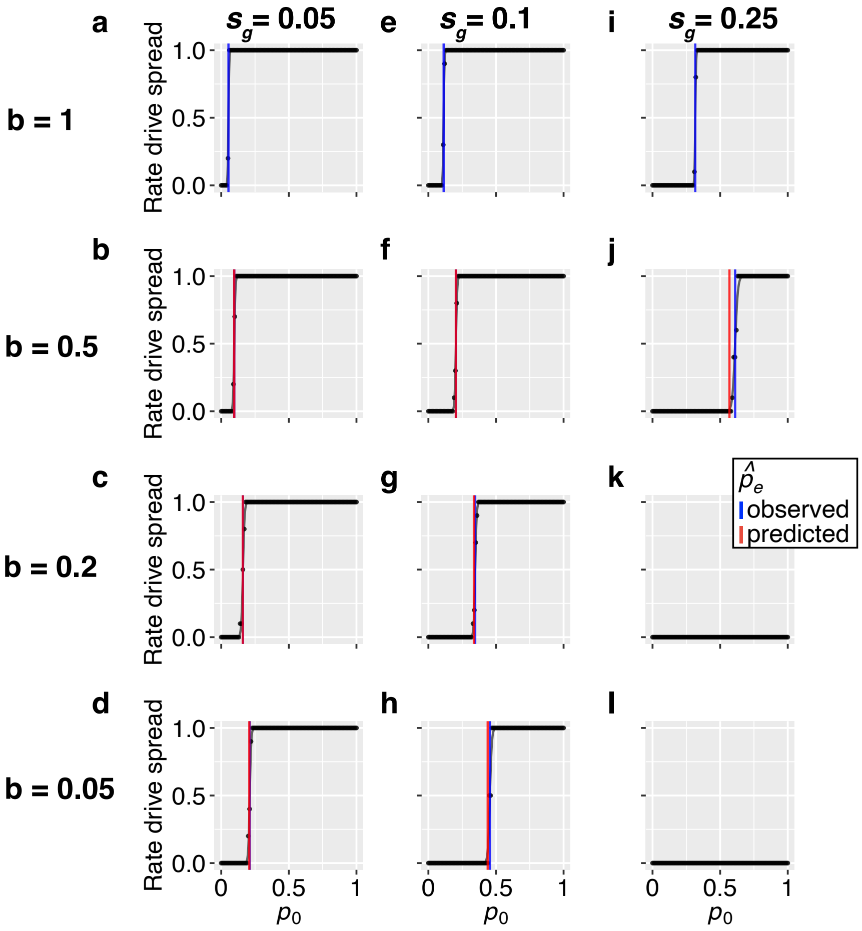


**Figure S14. Invasion ability of the ClvR modification drive under varying gamete viability fitness costs and baseline germination rates.** Figure design follows Figure S13 but shows results for the ClvR drive.

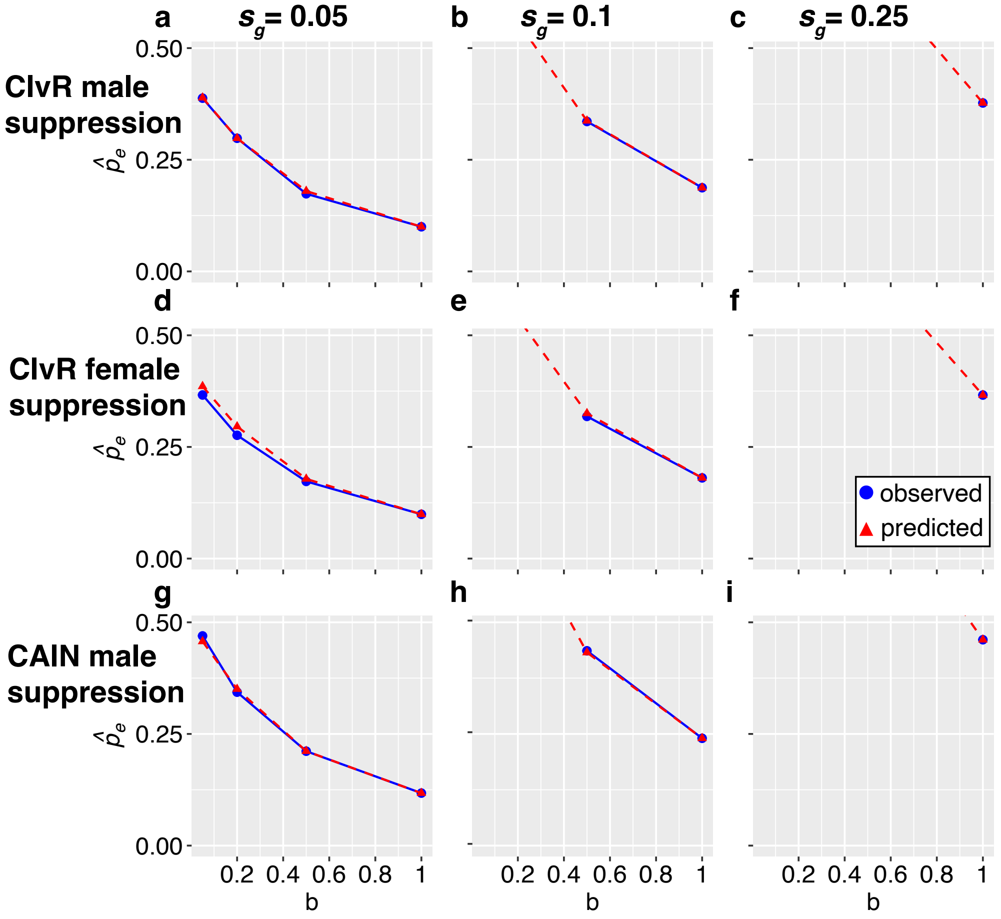


**Figure S15. Effective invasion thresholds of CAIN and ClvR suppression drives under gamete viability fitness costs.** Figure design follows Figure 6, except results are shown for suppression drives. For male suppression drives (ClvR in **a–c**, CAIN in **g–i**), the drive was introduced by converting a fraction $p_0$ of female plants to drive homozygotes, as male homozygotes for these drives are sterile. Conversely, for the ClvR female suppression drive (**d–f**), the drive was introduced by converting a fraction $p_0$ of male plants to drive homozygotes, as female homozygotes for this drive are sterile. Given an equal sex ratio, the maximum drive introduction frequency was 0.5. Absence of a point at $b = 0.05, 0.2, 0.5$, or 1 indicates either drive failure to spread (if blue) or a predicted effective invasion threshold exceeding 0.5 (if red). For further details on these drives' ability to spread, see Figures S16–S18.

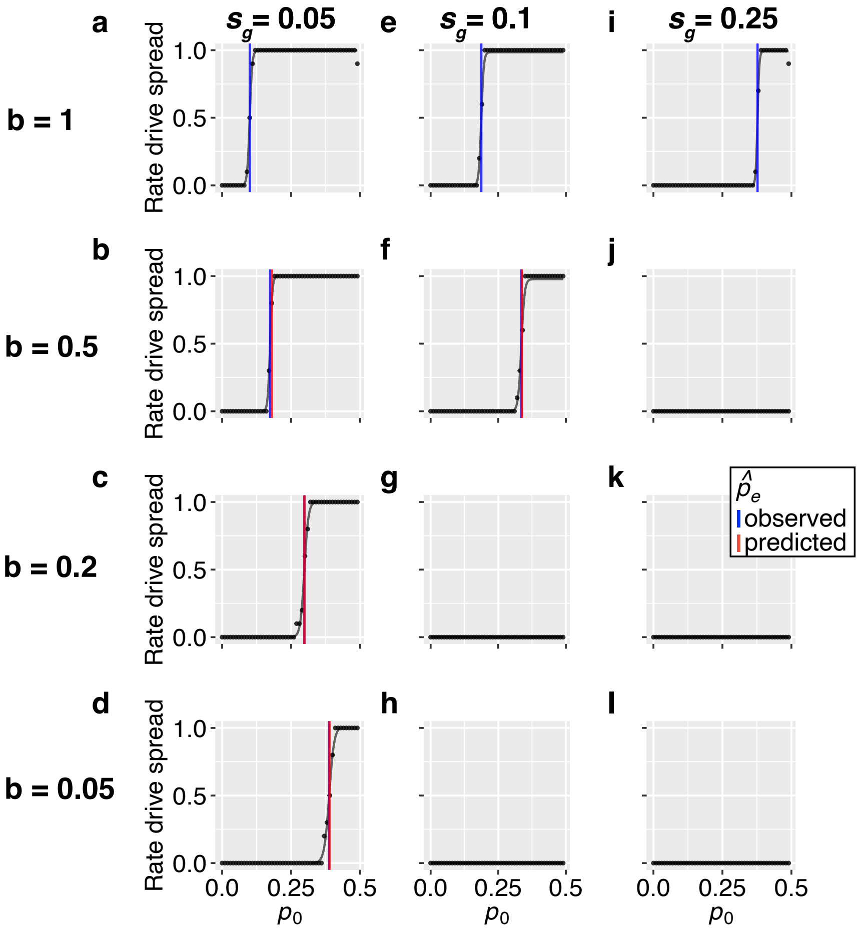


**Figure S16. Invasion ability of ClvR male suppression drives under varying gamete viability fitness costs and baseline germination rates.** Figure design follows Figure S13, except results are shown for ClvR male suppression. The drive was introduced by converting a fraction $p_0$ of female plants to drive homozygotes, as male drive homozygotes are sterile. With an equal sex ratio and a female-specific introduction, the maximum $p_0$ was 50%. Predicted effective invasion thresholds exceeding 50% are therefore not shown. Drive spread is defined as the drive eliminating the population or reaching a final frequency that exceeds $p_0$. **a–d,** $s_g = 0.05$ and $b = 0.05, 0.2, 0.5, 1$. **e–h,** $s_g = 0.1$ and $b = 0.05, 0.2, 0.5, 1$. **i–l,** $s_g = 0.25$ and $b = 0.05, 0.2, 0.5, 1$.

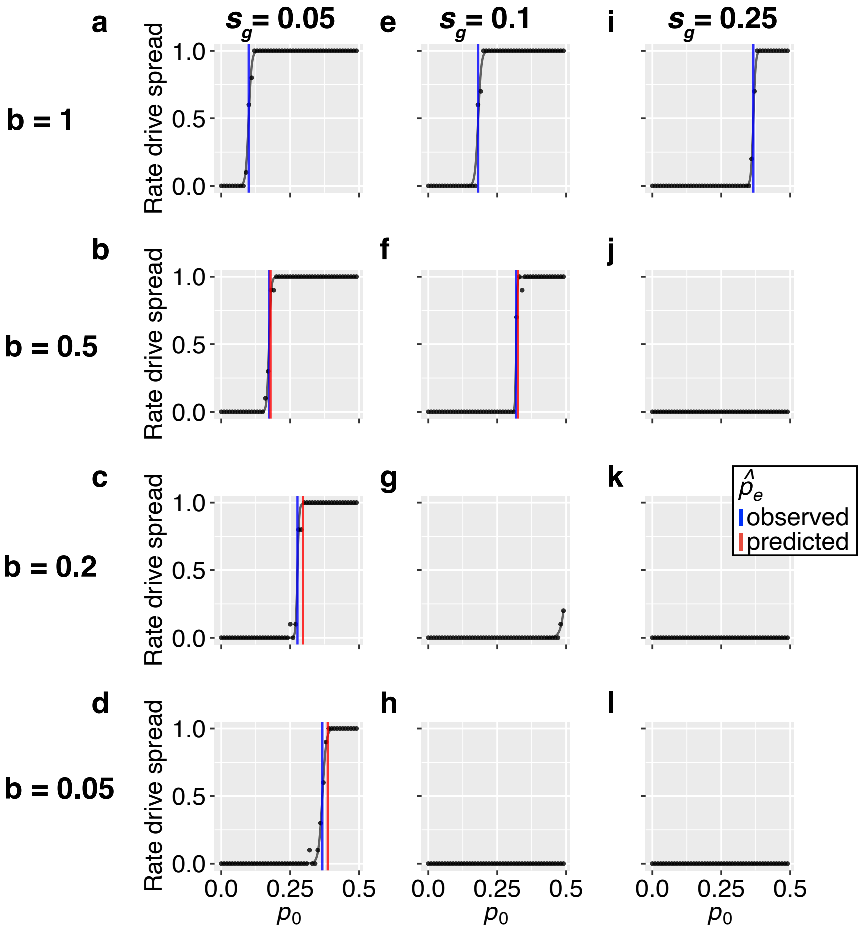


**Figure S17. Invasion ability of ClvR female suppression under varying gamete viability fitness costs and baseline germination rates.** Figure design follows Figure S13, except results are shown for ClvR female suppression. The drive was introduced by converting a fraction $p_0$ of male plants to drive homozygotes, as female drive homozygotes are sterile. Given a male-specific release and an equal sex ratio, the maximum $p_0$ was 0.5. Thus, predicted effective thresholds exceeding 0.5 are not shown. Drive spread is defined as the drive eliminating the population or reaching a final frequency that exceeds $p_0$.

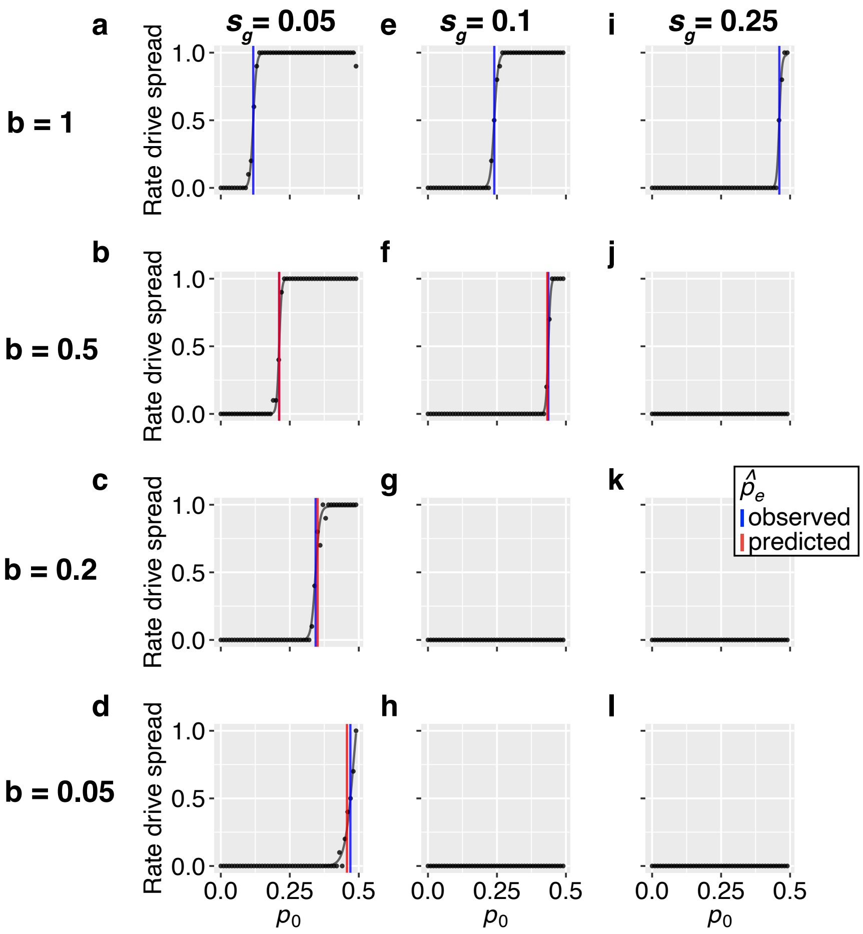


**Figure S18. Invasion ability of CAIN male suppression drives under varying gamete viability fitness costs and baseline germination rates.** Figure design follows Figure S13 but shows results for the CAIN male suppression drive. With a female-specific release and an equal sex ratio in the population, the maximum $p_0$ was 0.5. Predicted effective invasion thresholds exceeding 0.5 are therefore not shown. Drive spread is defined as the drive eliminating the population or reaching a final frequency that exceeds $p_0$.

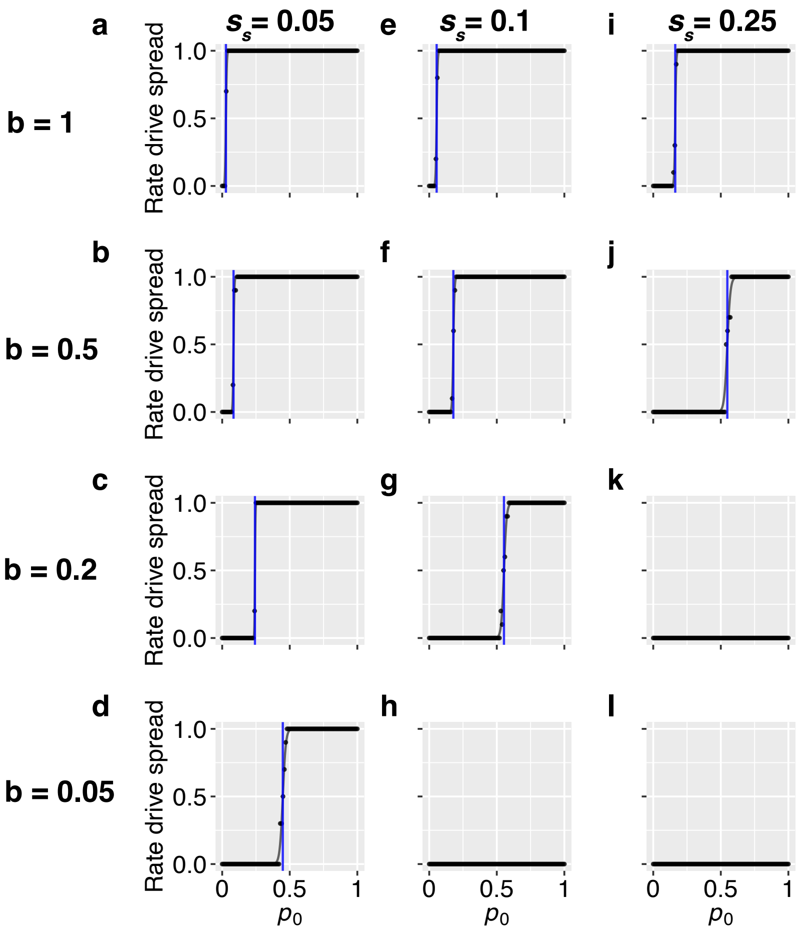


**Figure S19. Invasion ability of the CAIN modification drive under varying seed survival fitness costs and baseline germination rates.** For a given baseline germination rate ($b$) and drive fitness cost in seed survival ($s_s$), the drive was introduced by converting a fraction $p_0$ of plants to drive homozygotes. This drive fitness cost acts to reduce the baseline survival rate ($d$) of drive-carrying seeds and is assumed to be codominant, such that drive homozygous seeds have a baseline survival rate of $d - s_s$, and drive heterozygous seeds have a baseline survival rate of $d - s_s/2$. All other parameters were set to default values (Table S2). Each point shows the fraction of replicates (out of 10 total) in which the drive spread from a given introduction frequency ($p_0$). Drive spread is defined as reaching a final drive frequency that exceeds $p_0$. Logistic curves were fit using the `drc` R package [3]; the blue line indicates the observed effective invasion threshold, corresponding to the introduction frequency at which the drive spreads in 50% of replicates (where the logistic curve intersects with $y = 0.5$). **a–d,** $s_g = 0.05$ and $b = 0.05, 0.2, 0.5, 1$. **e–h,** $s_g = 0.1$ and $b = 0.05, 0.2, 0.5, 1$. **i–l,** $s_g = 0.25$ and $b = 0.05, 0.2, 0.5, 1$.

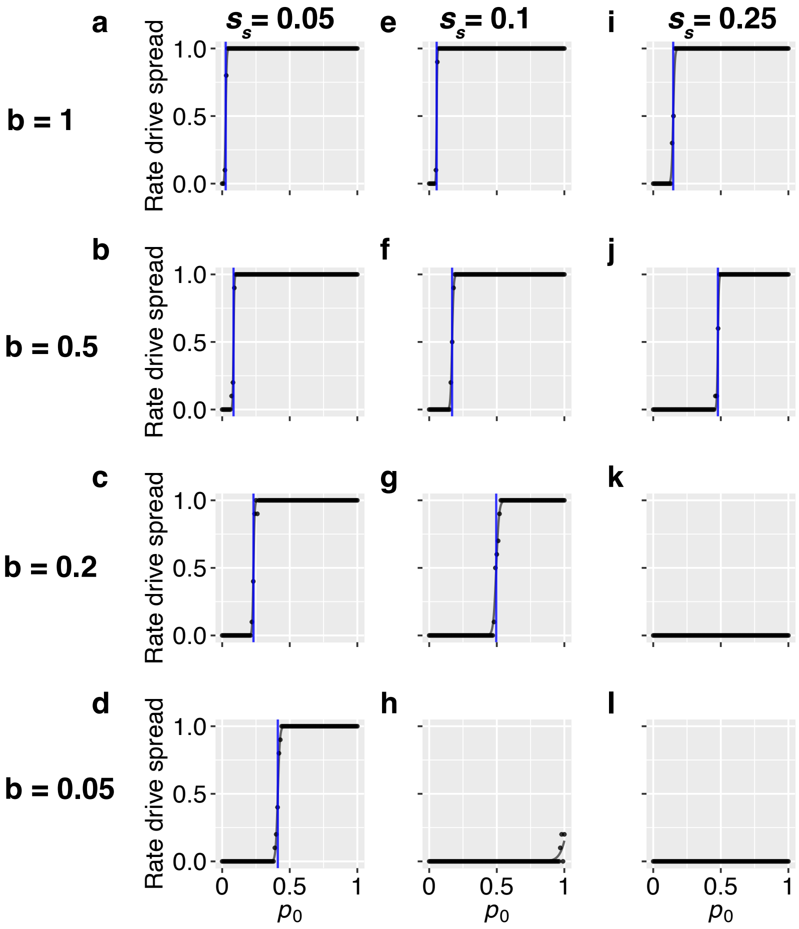


**Figure S20. Invasion ability of the ClvR modification drive under varying seed survival fitness costs and baseline germination rates.** Figure design follows Figure S19 but shows results for the ClvR drive.

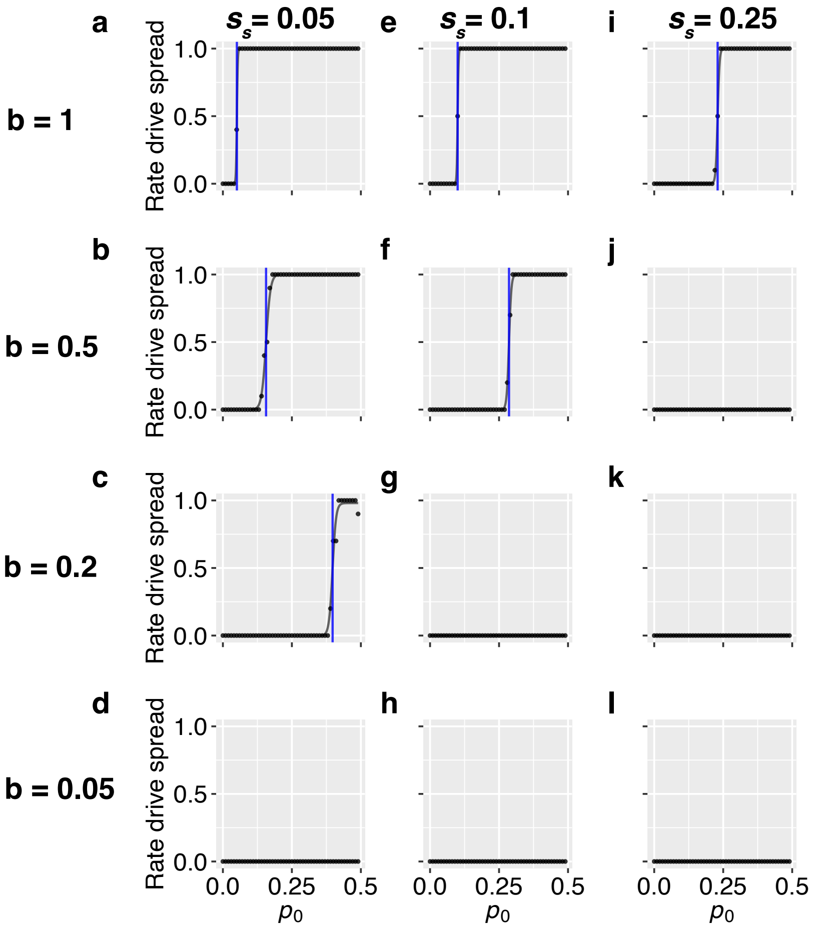


**Figure S21. Invasion ability of ClvR male suppression under varying seed survival fitness costs and baseline germination rates.** Figure design follows Figure S19, except results are shown for ClvR male suppression. The drive was introduced by converting a fraction $p_0$ of female plants to drive homozygotes, as male drive homozygotes are sterile. With a female-specific release and an equal sex ratio, the maximum $p_0$ was 0.5. Drive spread is defined here as the drive eliminating the population or reaching a final frequency that exceeds $p_0$.

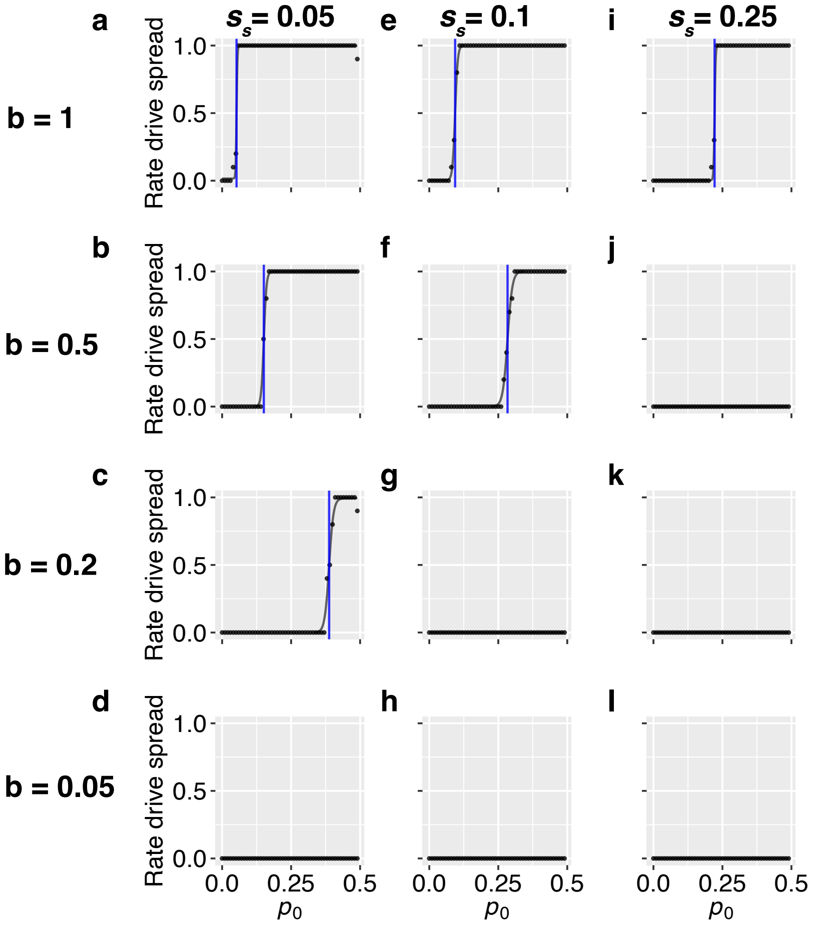


**Figure S22. Invasion ability of ClvR female suppression under varying seed survival fitness costs and baseline germination rates.** Figure design follows Figure S21, except the drive was introduced by converting a fraction of male plants to drive homozygotes, since female homozygotes for this drive are sterile. Given a male-specific release and an equal sex ratio in the population, the maximum $p_0$ was 0.5. Drive spread is defined as the drive eliminating the population or reaching a final frequency that exceeds $p_0$.

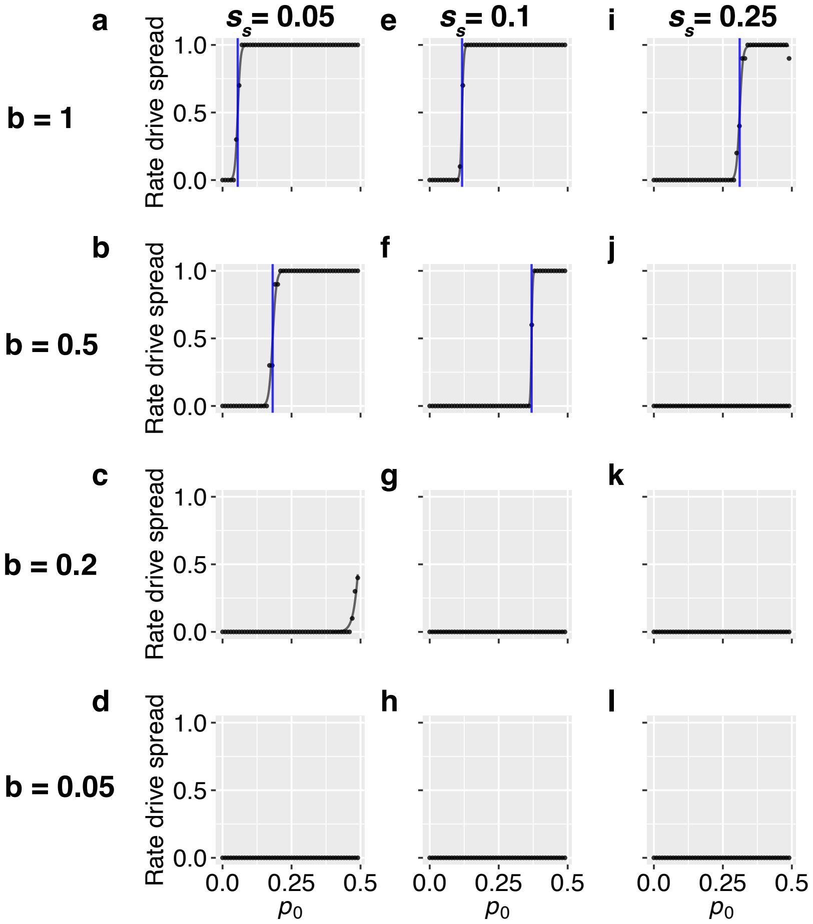


**Figure S23. Invasion ability of CAIN male suppression under varying seed survival fitness costs and baseline germination rates.** Figure design follows Figure S21, except results are shown for CAIN male suppression. With a female-specific release and an equal sex ratio, the maximum $p_0$ was 0.5. Drive spread is defined here as the drive eliminating the population or reaching a final frequency that exceeds $p_0$.

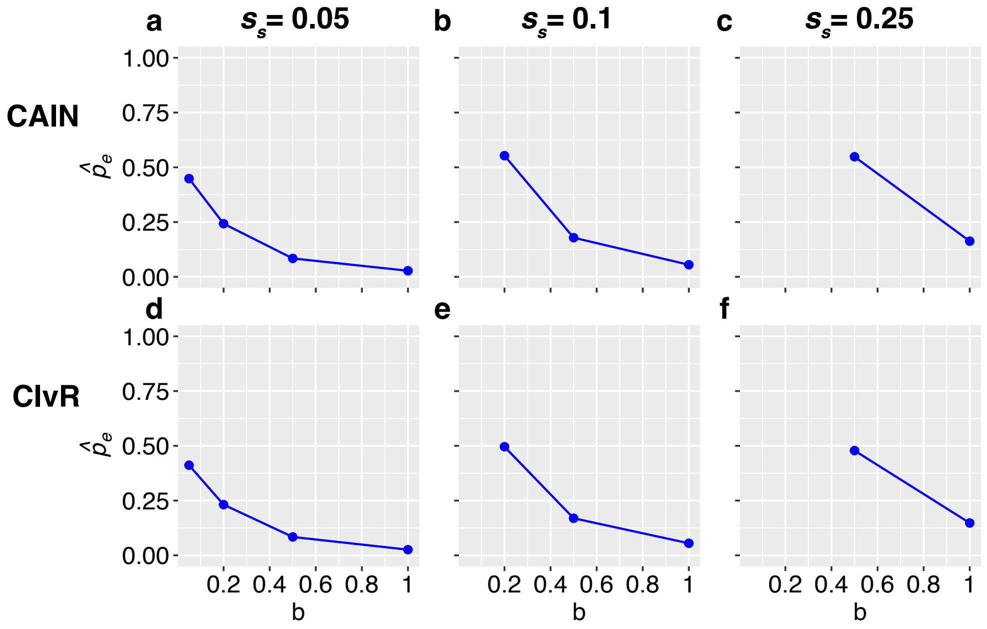


**Figure S24. Effective invasion thresholds of CAIN and ClvR modification drives under seed survival fitness costs.** Baseline germination rate ($b$) and drive fitness cost on seed survival ($s_s$) were varied, with all other parameters set to default values (Table S2). The fitness cost $s_s$ acts reduces the baseline survival rate ($d$) of drive-carrying seeds and is assumed codominant, such that drive homozygous seeds have a baseline survival rate $d - s_s$ and drive heterozygous seeds have a baseline survival rate $d - s_s/2$. Blue points show the observed effective invasion threshold, defined as the minimum introduction frequency above which the drive spreads in more than 50% of replicates (see Figures S19 and S20). **a–c,** Observed effective invasion thresholds for CAIN at $s_s = 0.05, 0.1, 0.25$ and $b = 0.05, 0.2, 0.5, 1$. Absence of a point at these $b$ values indicates that the drive consistently failed to spread. **d–f,** Same as **a–c** but for ClvR.

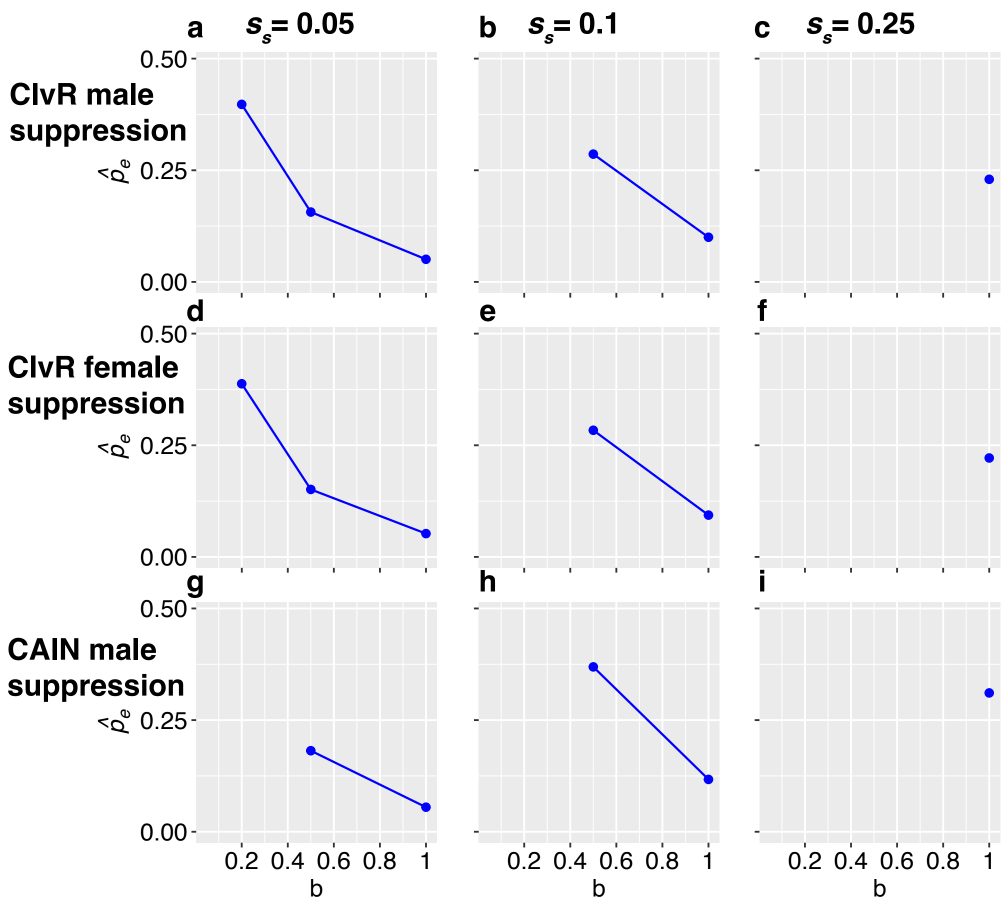


**Figure S25. Effective invasion thresholds of CAIN and ClvR suppression drives under seed survival fitness costs.** Figure design follows Figure S24. For male suppression drives (ClvR in **a–c**, CAIN in **g–i**), the drive was introduced by converting a fraction of female plants to drive homozygotes, since male homozygotes are sterile. Conversely, for the ClvR female suppression drive (**d–f**), the drive was introduced by converting a fraction of male plants to drive homozygotes, since female homozygotes are sterile. Given sex-specific releases and an equal sex ratio, the maximum introduction frequency was 50%. Blue points show the observed effective invasion threshold, defined as the introduction frequency above which the drive spreads in more than 50% of replicates (see Figures S21–S23). **a–c,** Observed effective invasion thresholds for ClvR male suppression at $s_s = 0.05, 0.1, 0.25$ and $b = 0.05, 0.2, 0.5, 1$. Absence of a point at these $b$ values indicates that the drive consistently failed to spread. **d–f,** Same as **a–c** but for ClvR female suppression. **g–i,** Same as **a–c** but for CAIN male suppression.
